# A universal pocket in Fatty acyl-AMP ligases ensures redirection of fatty acid pool away from Coenzyme A-based activation

**DOI:** 10.1101/2021.05.04.442589

**Authors:** Gajanan S. Patil, Priyadarshan Kinatukara, Sudipta Mondal, Sakshi Shambhavi, Ketan D. Patel, Surabhi Pramanik, Noopur Dubey, Subhash Narasimhan, Murali Krishna Madduri, Biswajit Pal, Rajesh S. Gokhale, Rajan Sankaranarayanan

## Abstract

Fatty acyl-AMP ligases (FAALs) channelize fatty acids towards biosynthesis of virulent lipids in mycobacteria and other pharmaceutically or ecologically important polyketides and lipopeptides in other microbes. They do so by bypassing the ubiquitous coenzyme A-dependent activation and rely on the acyl carrier protein-tethered 4’-phosphopantetheine (*holo*-ACP). The molecular basis of how FAALs strictly reject chemically identical and abundant acceptors like coenzyme A (CoA) and accept *holo*-ACP unlike other members of the ANL superfamily remains elusive. We show FAALs have plugged the promiscuous canonical CoA-binding pockets and utilize highly selective alternative binding sites. These alternative pockets can distinguish adenosine 3’, 5’-bisphosphate-containing CoA from *holo*-ACP and thus FAALs can distinguish between CoA and *holo*-ACP. These exclusive features helped identify the omnipresence of FAAL-like proteins and their emergence in plants, fungi, and animals with unconventional domain organisations. The universal distribution of FAALs suggests they are parallelly evolved with FACLs for ensuring a CoA-independent activation and redirection of fatty acids towards lipidic metabolites.

## INTRODUCTION

The ANL superfamily includes enzymes such as the Acyl/Aryl-CoA ligases (ACS or FACLs), Adenylation domains (A-domains) and Luciferases along with the recently identified Fatty acyl-AMP ligases (FAALs). These enzymes are involved in the production of both primary metabolites such as acyl-CoA and secondary metabolites such as antibiotics ^1^, complex lipids ^2^, cyclic peptides ^3^ and lipopeptides ^4,5^. Basic metabolic pathways such as β-oxidation, membrane biogenesis, post-translational modifications etc., use primary metabolites such as acyl-CoA. The secondary metabolites such as complex lipids that function as virulent molecules in *Mycobacteria* and bioactive molecules in several microbes that help tide over unfavourable conditions and establish themselves in their niches. Such diverse metabolites are produced by the members of the ANL superfamily through a two-step catalytic mechanism. It begins with the activation of carboxylate-moiety of substrates such as fatty acids or amino acids by ATP hydrolysis and finally transferring it to an acceptor such as CoA or *holo*-ACP. Multiple structural and biochemical studies show that members of the superfamily such as FACLs and A-domains employ a common pocket for the chemically identical CoA and the 4’-PPant moieties attached to the *holo*-ACP, respectively for the final transfer. It was later demonstrated that the A-domains can cross-react with CoA to form aminoacyl-CoA ^6^, which points to the liabilities of utilizing a common pocket for binding chemically identical moieties. Infidelity towards the final acceptor has now been noted in different classes of ANL superfamily members where Luciferases are shown to catalyse fatty acyl-CoA formation ^7^ and FACLs producing bioluminescence with molecular oxygen ^8^. While fatty acid/amino acid substrate promiscuity is well studied and exploited in combinatorial biosynthesis of bioactive molecules, the origin and basis of acceptor promiscuity is relatively less understood.

FAALs are atypical enzyme systems of the ANL superfamily as they completely lack acceptor promiscuity, where they transfer the activated fatty acyl-AMP to the 4’-PPant of *holo*-ACP ^9^ but not CoA. FAALs rejecting the small, diffusible, and abundant CoA while accepting the ACP-tethered to a 4’-PPant moiety (Supplementary Figure-1) is puzzling as they are chemically identical. In a previous study, it was proposed that the FAAL-specific insertion (FSI), an additional stretch of amino acids found only in the N-terminal domain of FAALs, prevents reaction with CoA^10^. However, the deletion or destabilization of the FSI failed to convert FAALs as efficient producers of acyl-CoA as there is only a weak ability to react with CoA ^10,11^. Moreover, it is also unclear how such a mechanism operates and distinguishes the two chemically identical acceptors, particularly when CoA is an abundant metabolite. These observations prompted us to hypothesize that FAALs have either evolved novel appendages or other modes for binding the acceptor to allow strict rejection of CoA.

In the present study, we have used structural, mutational, and biochemical analysis to identify the mechanistic basis of how FAALs can distinguish between near identical acceptors for the acyl transfer reaction. We show that, unlike other members of the superfamily, FAALs achieve acceptor fidelity by avoiding the usage of a promiscuous canonical CoA-binding pocket and utilizing a discriminatory pocket that is distinct from the canonical CoA-binding pocket. Loss- and gain-of-function mutations were generated by identifying the structural determinants that nullify the canonical CoA-binding pocket. Interestingly, we found that the non-functional canonical CoA-binding pocket and the unique discriminatory alternative pocket are unique features of FAALs which is also conserved in all forms of life including plant, fungi, and animals. The identification of such a conserved rejection mechanism across organisms has larger implications in determining the redirection of cellular flux of fatty acids towards synthesis of diverse metabolites across organisms.

## RESULTS AND DISCUSSION

### The promiscuous canonical CoA-binding pocket is inaccessible and redundant in FAALs

The structural features of a canonical CoA-binding pocket that allow the proper recognition of CoA/4’-PPant in different ANL superfamily members was compared and contrasted with the analogous structural positions in FAALs. A comprehensive analysis of the canonical CoA-binding pocket in the 26 structures (59 protomers) of the CoA/4’-PPant-bound ANL superfamily members (Supplementary Table-1) revealed important aspects of CoA/4’-PPant recognition. It is observed that ligand interacts with the protein through three categories of interactions but none of the structures show the CoA/4’-PPant ligands bound in identical positions or orientations. The three categories of interactions are; (i) the hydrogen bonds mediated by the A8-motif of the C-terminal domain (ii) the water-mediated contacts through (Supplementary Figure-2b) the N-terminal helices H10 and H14 (notation based on *Escherichia coli* FAAL abbreviated as *Ec*FAAL; PDB ID: 3PBK) ^12^ and (iii) the interactions with the phosphates ^13-20^ of the adenosine 3’,5’-bisphosphate moiety through of the positively charged residues (Arg/Lys) (Supplementary Figure-3a) in the loop connecting H23 and β26 (notation based on *Ec*FAAL).

A comparative analysis of the structurally analogous CoA-binding pocket of FAALs (11 structures constituting 23 protomers) ^10-12,21,22^ sheds light on why FAALs cannot accept CoA. The analysis reveals an absence of selection from the positively charged residues (Arg/Lys) known to assist in CoA binding in the other members of the ANL superfamily (Figure-1a). In addition, the CoA-binding pocket of FAALs shows the presence of bulky residues F279, W224 and M231 (in *Ec*FAAL) at the base and a unique FAAL-specific helix (FSH) (T252-R258 in *Ec*FAAL) at the entrance of the pocket (Figure-1b). These elements are highly conserved in other FAALs (Figure-1c) and restrict the space available for the incoming 4’-PPant arm of both CoA and *holo*-ACP. The superposition of CoA- and 4’-PPant-bound structures with FAALs revealed that the overcrowded pocket along with the FSH at the entrance of the pocket is unlikely to accommodate both CoA or 4’-PPant of *holo*-ACP and hinder their entry (Figure-1b). There are at least seven atoms of the N-terminal domain of FAALs at a distance less than 2.5 Å from the atoms of CoA in its various conformations found in the different CoA-bound FACL structures. On the contrary, the N-terminal domain of FACLs shows only one atom at 2.5 Å from the multiple CoA conformations (Supplementary table-II). The distance-based assessment clearly indicates the potential clashes an incoming CoA would face in the canonical pocket of FAALs. Moreover, the absence of positively charged residues will naturally be unfavourable for CoA from being appropriately oriented in the pocket. These observations lead to the hypothesis that the canonical CoA-binding pocket is rendered non-functional because of the inaccessibility in FAALs.

**Figure-1:**
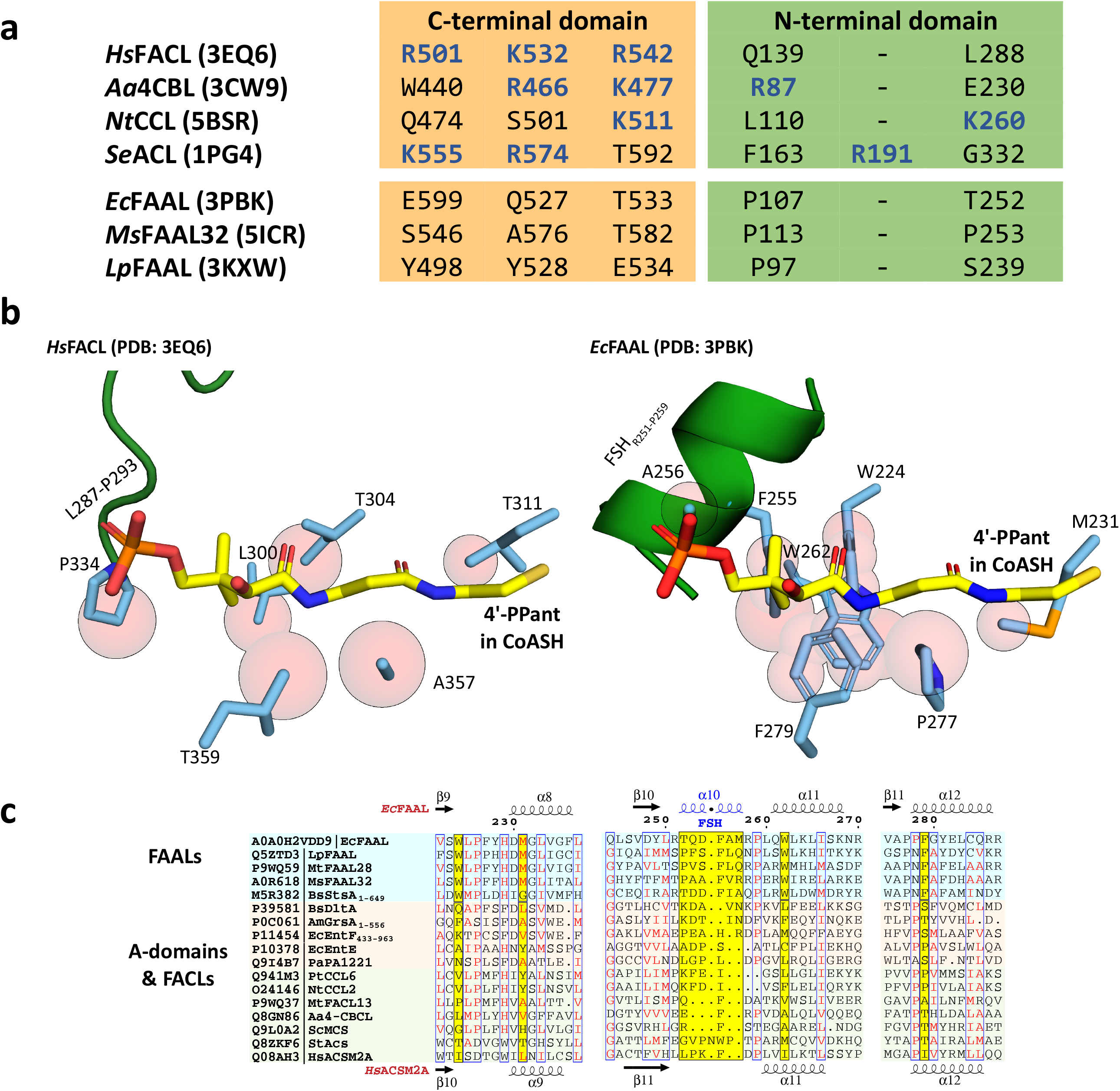
The presence of conserved negative selection elements and absence of positive selection elements in canonical CoA-binding pocket of FAALs can prevent CoA-binding: (**a**) The CoA-interacting residues in FACLs (*Hs*FACL, *Aa*FACL, *Nt*FACL, *Se*FACL) and the structurally analogous residues in FAALs (*Ec*FAAL, *Ms*FAAL32 and *Lp*FAAL) are tabulated. The positively charged residues (blue) mainly are from the C-terminal domain (orange) and occasionally from the N-terminal domain (green). (**b**) The residues (cyan) in the vicinity (≤ 2.5 Å) of the bound CoA (yellow; *Hs*FACL, PDB: 3EQ6) are shown as van der Waal spheres (light red) to compare the canonical pocket in FAALs and FACLs. The adenosine-3’-5’-bisphosphate moiety of CoA is omitted in the representation for clarity. A unique FAAL-specific helix (FSH_R251-P259_) at the entry of the canonical pocket shown in cartoon representation (green) is replaced by a loop (L287-P293) in FACLs. (**c**) The canonical pocket obstructing features seen in FAALs (light cyan) are highlighted in yellow in the structure-based sequence alignment and compared to other representative members of the ANL superfamily (FACLs in light green and A-domains in light orange). The secondary structures of *Ec*FAAL (PDB: 3PBK) and *Hs*FACL (PDB: 3EQ6) are marked at the top and bottom of the alignment, respectively.

### Resurrecting the canonical CoA pocket enables gain of function in FAALs

The structural analysis was used to design mutations in FAALs and FACLs where the bulkier residues of the canonical CoA-binding pocket of FAALs were mutated to smaller residues to induce “gain of function” (Figure-2a, 2b). Likewise, the smaller residues in the canonical CoA-binding pocket of FACLs were substituted with the corresponding conserved bulkier residues present in FAALs to measure the “loss of function” (Figure-2d, 2e). Individual mutations reducing the size of residues in the canonical CoA-binding pocket resulted in the production of acyl-CoA or “gain of function” in FAALs that otherwise does not make any acyl-CoA (Figure-2c). A considerable amount of the total acyl-AMP formed was converted to acyl-CoA, ∼80% in the case of *Ms*FAAL32_Δ254-257_ and ∼60% in the case of *Rs*FAAL_Δ240-243_, when the FSH segment was deleted as compared to their wild-type proteins, respectively. Some of the single point mutants such as A253F in *Mt*FACL13 show reduced production of acyl-CoA by ∼80%, while mutations in combinations such as A276F/A232M of *Af*FACL reduce the turnover of acyl-AMP to acyl-CoA by 98% as compared to wild-type (100%) (Figure-2f). These observations clearly indicate that the size of residues in the canonical CoA-binding pocket dictate the ability to accommodate CoA and hence the ability to facilitate the thioesterification reaction with CoA.

**Figure-2:**
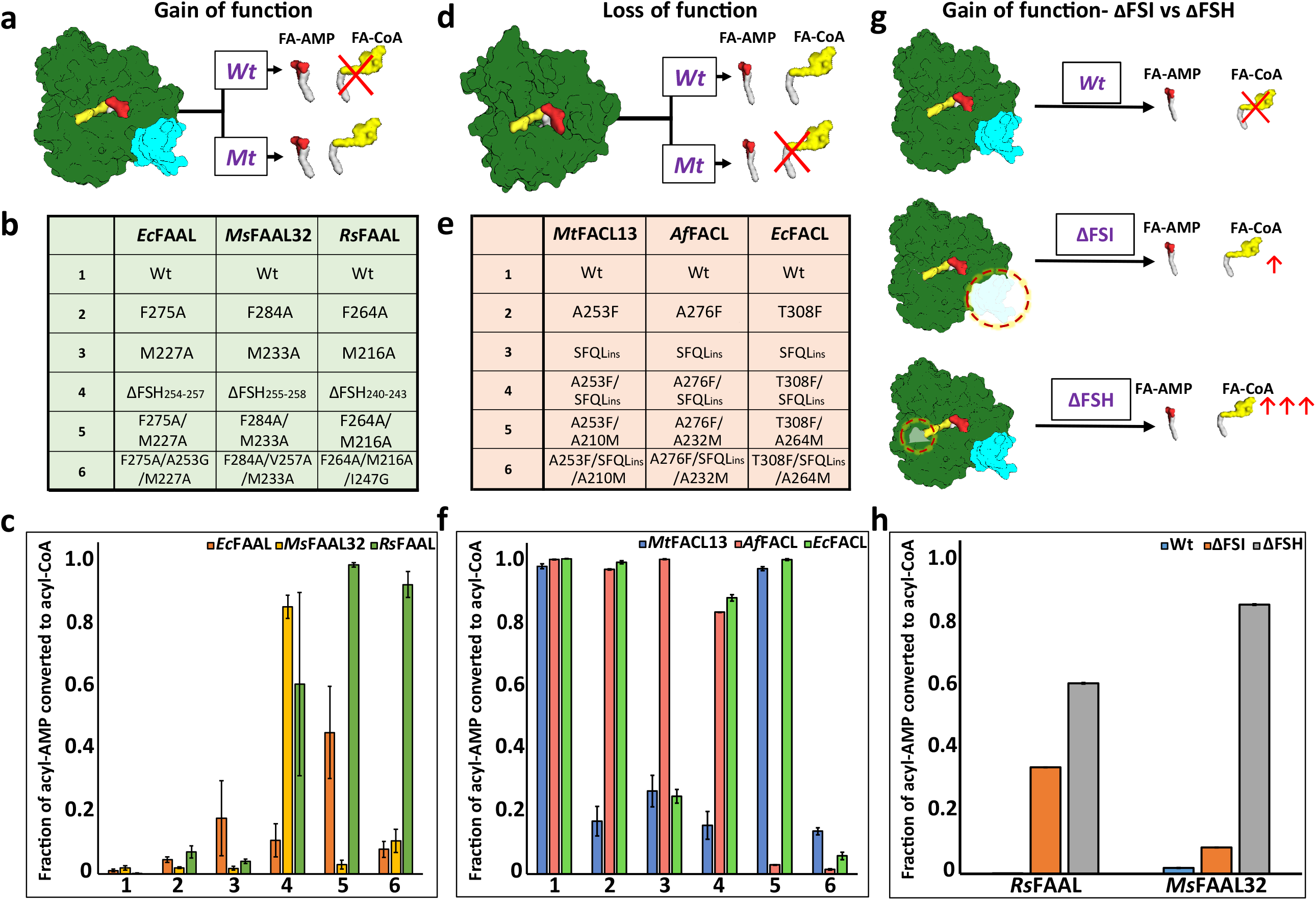
Biochemical analysis of “gain of function” mutants of FAALs and “loss of function” mutants of FACLs: (**a**) The biochemical activity of a wild-type (Wt) FAAL as compared to its mutants (Mt) is schematically represented. (**b**) The various “gain of function” mutations of FAALs (*Ec*FAAL, *Ms*FAAL32 & *Rs*FAAL) are tabulated as the following: Row-1 = Wild-type protein; Rows-2 and -3 = single point mutations; row-4 = deletion of FSH and rows-5 and -6 = combinations of these mutations. (**c**) The fraction of acyl-AMP converted to acyl-CoA by wild type and various mutants (represented as numbers on the x-axis according to Figure-2b) of FAALs is represented as a bar graph along with standard error of the mean. (**d**) The biochemical activity of a wild-type (Wt) FACL as compared to its mutants (Mt) is schematically represented. (**e**) The various “loss of function” mutations generated in FACLs (*Mt*FACL, *Af*FACL & *Ec*FACL) are tabulated as the following: Row-1 = wild-type protein; row-2 = single point mutations; row-3 = insertion of FSH; and rows-4, -5 and -6 = combinations of these mutations. (**f**) The fraction of acyl-AMP converted to acyl-CoA by wild type and various mutants (represented as numbers on the x-axis according to Figure-2e) of FACLs is represented as a bar graph along with standard error of the mean. (**g**) A comparison of the gain of function from ΔFSH mutation with ΔFSI mutation in FAALs is schematically represented. (**h**) The fraction of acyl-AMP converted to acyl-CoA by wild type, ΔFSH and ΔFSI mutants of *Rs*FAAL and *Ms*FAAL32 (represented on the x-axis) is represented as a bar graph along with standard error of the mean.

It was previously identified that deletion of FAAL-specific insertion (ΔFSI) can lead to gain of function in FAALs ^10^. Current results indicate that mutations in the canonical pocket alone is sufficient to introduce CoA production ability in FAALs. A comparison of acyl-CoA production of the ΔFSI with FSH deletion mutant (ΔFSH) (Figure-2g) reveals that *Rs*FAAL_ΔFSH_ produces 2-fold excess acyl-CoA as compared to *Rs*FAAL_ΔFSI_ while *Ms*FAAL32_ΔFSH_ produces 10-fold excess acyl-CoA as compared to *Ms*FAAL32_ΔFSI_ (Figure-2h). Therefore, it can be concluded that the FSH and other CoA-rejection elements around the canonical CoA-binding pocket significantly deter acyl-CoA production, which supersedes the inhibition of acyl-CoA production by FSI in FAALs. Overall, these observations lead to the conclusion that the available space in the canonical CoA-binding pocket of FAALs is very small to accommodate even the 4’-PPant moiety. Mutations that increase the pocket space enhance the acyl-CoA production in FAALs, while the opposite is observed in the case of FACLs. Therefore, the bulky residues in the putative canonical CoA-binding pocket of FAALs itself acts as a negative selection gate to sterically exclude CoA. The rejection mechanism operational at the canonical CoA-binding pocket in FAALs is relatively more effective in rejecting CoA than the FSI-based rejection.

### Identification of an alternative 4’-PPant-binding pocket in FAALs

The steric occlusion of the 4’-PPant binding in canonical pocket would not only result in prevention of CoA-binding but also restrict the access to the 4’-PPant tethered *holo*-ACP. Previous studies have highlighted that FAALs work in coordination with *holo*-ACP’s, stand-alone or fused to polyketide synthase (PKS) or nonribosomal peptide synthetase (NRPS) modules, to produce various bioactive molecules such as complex lipids of *Mycobacterium*, lipopeptides of *Ralstonia* and other cyanobacterial species. Since the canonical pocket is rendered inaccessible, it immediately prompts the question ‘how then FAALs are able to transfer the activated fatty acids to the 4’-PPant arm of the *holo*-ACP? We, therefore, sought to identify if FAALs have evolved an alternative mechanism to accommodate the 4’-PPant arm to allow binding and subsequent catalysis. Analysis of the crystal structures of the N-terminal domains of FAALs using various pocket search algorithms such as MOLE 2.0 ^23^, DOGSiteScorer ^24,25^, PyVOL ^26^, KVFinder ^27^ and POCASA ^28^ helped us in identifying a novel cavity in the N-terminal domain of FAALs (Figure-3a) but not identifiable in any of the known crystal structures of FACLs.

**Figure-3:**
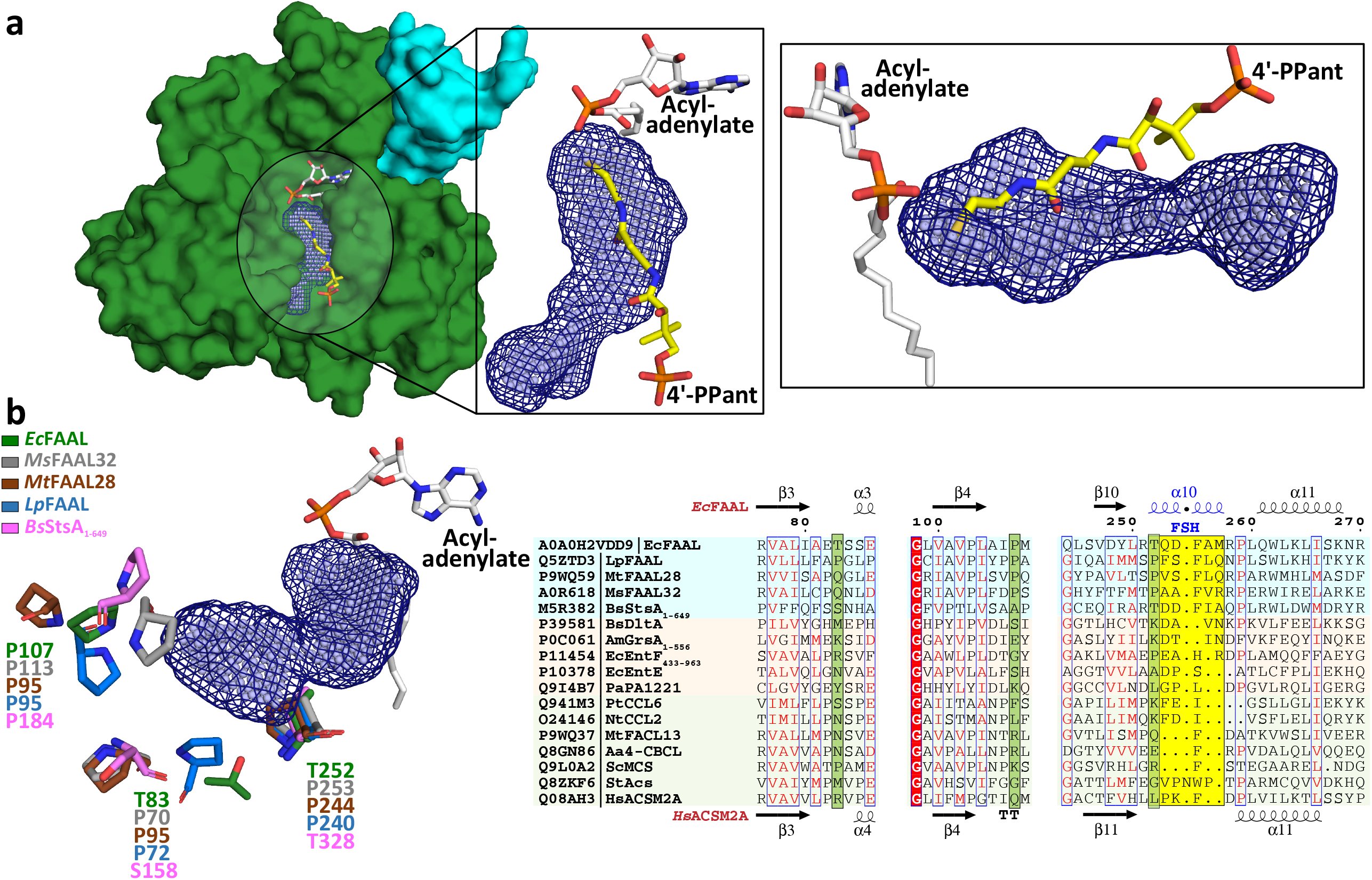
FAALs have an alternative pocket in the N-terminal domain distinct from the canonical CoA-binding pocket and lined by conserved prolines: (**a**) The analysis of N-terminal domain of *Ec*FAAL (PDB: 3PBK) (green; surface representation) with pocket finding algorithms identified a new pocket (dark blue; mesh representation and light blue; spheres representation). Also shown are the FAAL-specific insertion (cyan), an *Ec*FAAL-bound lauryl-adenylate (grey; sticks representation) and the 4’-PPant moiety (yellow; sticks representation) of a CoA bound to *Hs*FACL (PDB: 3EQ6) in the canonical pocket. The inset shows two different orientations (top-view and lateral-view) of the distinct alternative pocket poised against the acyl-adenylate and compared to the 4’-PPant of a CoA in canonical pocket. (**b**) Various residues (stick representation) of FAALs residing at the entrance of the unique pocket (dark blue; mesh representation and light blue; spheres representation) are identified through structural comparison. A structure-based sequence alignment of these residues (highlighted in pale green) with other representative members of the ANL superfamily reveals that FAALs have a higher frequency of prolines (occasionally Thr/Ser) than other members of the superfamily.

The entrance of the distinct tunnel is on the N-terminal side of the FSH, while the canonical pocket to accommodate CoA is on the C-terminal side of the FSH. The approach towards the active site in both cases is not on the same plane, but they coincide near the active site near the β-alanine of the 4’-PPant. The longest length along this pocket is aligned at ∼25° to the canonical CoA-binding pocket. The canonical pocket is mainly formed after the rotation of the C-terminal domain in the thioesterification state (T-state) bringing the A8-motif near the active site. The space generated between the A8-motif (from the C-terminal domain) and the subdomain-B of the N-terminal domain constitute the canonical CoA-binding pocket. In contrast, the newly identified pocket is the space between the loops in subdomain-A and the FSH region of subdomain-B of FAALs. These loop regions are highly variable in length but rich in prolines (occasionally threonine or serine) some of which are conserved in FAALs such as P226 and P107 in *Ec*FAAL (Figure-3b), while the FSH has a unique secondary structure characteristic of FAALs. The structurally analogous sites in FACLs show a high degree of variability with occasional prolines but the frequency of prolines at the indicated positions is poor and often replaced by Asn/Leu (Supplementary Figure-4). The ability to consistently identify the tunnel with regions enriched in prolines around the opening led to the proposition that this alternative pocket is perhaps a universal attribute of FAALs that has evolved to accommodate the 4’-PPant arm.

### The alternative pocket in FAALs is functional and accepts a 4’-PPant-tethered ACP

The ability of the alternative pocket to accommodate 4’-PPant arm tethered to ACP was tested using structure-guided mutagenesis. Bulkier residues (Phe/Arg) were introduced at the entrance of the tunnel that could potentially block the accessibility of the pocket for the incoming 4’-PPant arm. The biochemical analysis of these mutants requires an assay system to monitor the transfer of acyl-AMP to the 4’-PPant arm of *holo*-ACP. Typically, such acyl-transfers are assessed using SDS-PAGE ^9,29^ or conformationally sensitive Urea-PAGE (CS-PAGE) ^30^. The ACPs that accept the acyl-chain from FAALs presented multiple complications such as poor conversion from *apo*-ACP to *holo*-ACP (Supplementary Figure-5a) and lack of separation on a CS-PAGE (Supplementary Figure-5b). Therefore, the CS-PAGE assay was modified for enhanced detection of the FAAL-dependent acyl-transfer on *holo*-ACP for the first time using the radio-labelled fatty acids. We tested the efficacy of the modified radio-CS-PAGE assay to probe three pairs of FAAL-ACP systems from diverse organisms (Figure-4a), viz FAAL-ACP pairs from *E. coli* (*Ec*FAAL-*Ec*ACP), *Myxococcus xanthus* (*Mx*FAAL-*Mx*ACP) and *Ralstonia solanacearum* (*Rs*FAAL-*Rs*ACP). The appearance of bright bands on the radio-CS-PAGE indicates that the radiolabelled fatty acid tethered to ACP, which is absent when *apo*-ACP is used, or ATP is omitted in the reaction. Thus, the assay system can enable the simultaneous probing of multiple FAAL-ACP pairs along with their mutations and facilitate similar studies in the future.

**Figure-4:**
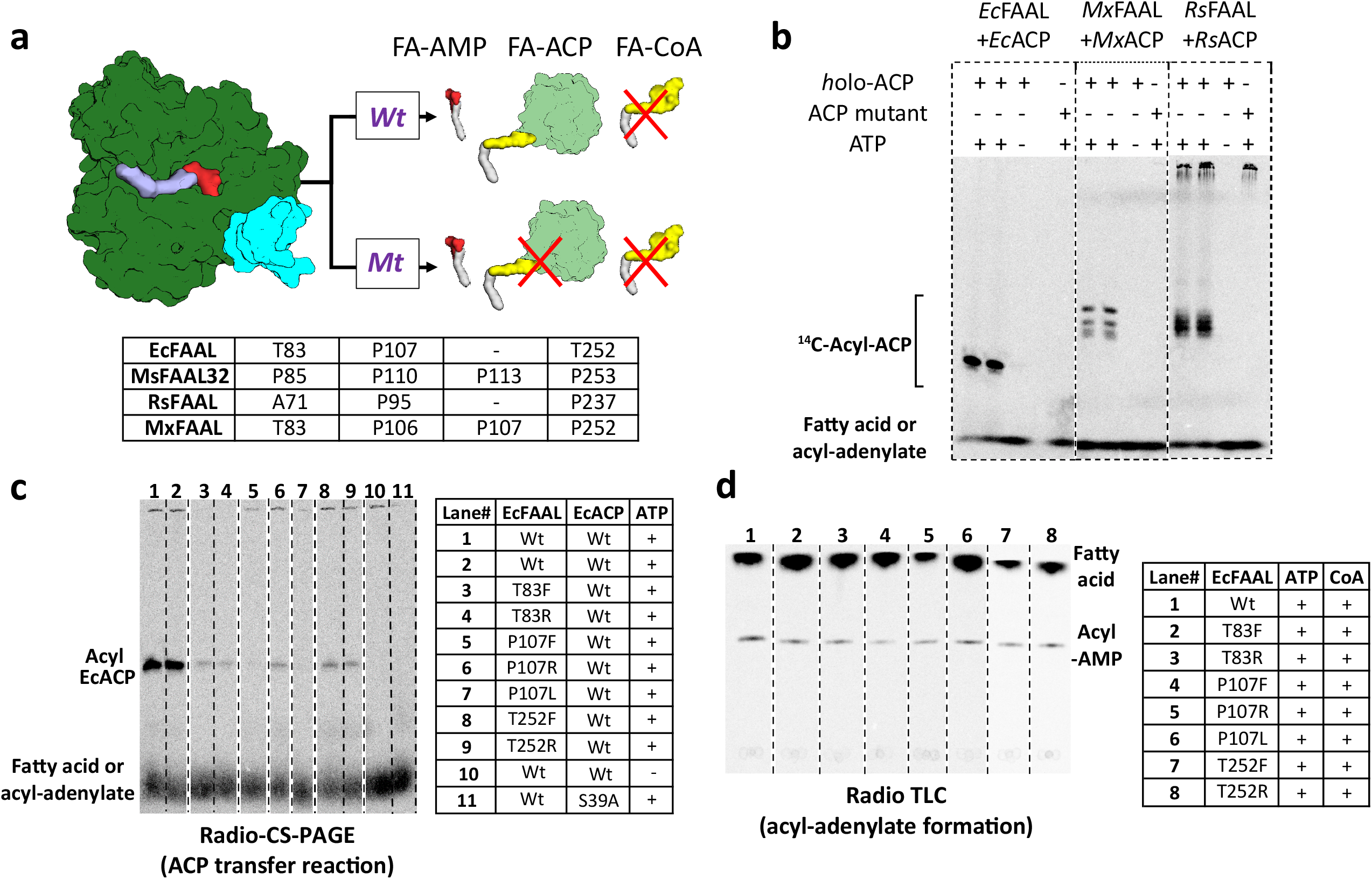
The alternative pocket identified in FAALs is a functional pocket that assists in catalysis by accommodating 4’-PPant tethered to ACP: (a) A schematic representation of the acyl-transfer function in a wild type (Wt) FAAL is compared to a mutant protein (Mt) that blocks the alternative pocket. (b) A representative radio-CS-PAGE gel shows the acyl-transfer reaction by wild type FAALs on their respective *holo*-ACP. Three representative FAALs (*Ec*FAAL, *Mx*FAAL and *Rs*FAAL) catalyse the transfer of radioactive acyl group on *holo*-ACP (observed as a bright band on the gel). The reaction product is not observed if ATP is not included in the reaction or when a mutant ACP (conserved serine mutated to Alanine) without the 4’-PPant arm is used. (c) The radio-CS-PAGE shows that mutations of the residues lining the alternative pocket to Arg/Phe show a reduced or negligible acyl-ACP formation. (d) The radio-TLC shows that loss of acyl-ACP formation is not due to affected acyl-adenylate formation and these mutants do not form any acyl-CoA.

The modified radio-CS-PAGE was then used to test if mutating prolines (occasionally a threonine) guarding the entrance of the pocket to bulkier residues can cause abrogation of the acyl-transfer activity (Figure-4c). It was found that a single point mutation, T83F or T83R, T252F or T252R and P107F or P107R in *Ec*FAAL can almost abrogate the acyl-transfer reaction on *holo*-*Ec*ACP (Figure-4c). It should be noted that these mutations did not affect the adenylation ability of the proteins (Figure-4d). The mutations in other FAAL-ACP pairs, *Mx*FAAL-*Mx*ACP and *Rs*FAAL-*Rs*ACP also resulted in abrogation of the acyl-transfer ability on their respective cognate *holo*-ACPs. An additional FAAL-PKS pair of *Ms*FAAL32-*Ms*PKS13_1-1042_ was also mutated and probed biochemically using the traditional radio-SDS-PAGE. The mutations of residues guarding the alternative pocket to bulkier residues in the *Ms*FAAL32-*Ms*PKS13_1-1042_ system also resulted in diminished acyl-transfer ability. These results indicate that blocking the entrance of the alternative pocket prevents the entry of the incoming 4’-PPant arm tethered to ACP. Therefore, in FAALs, the identified alternative pocket is fully functional and distinct from the non-functional canonical pocket. Such a unique pocket facilitates the entry of the 4’-PPant arm of the ACP to approach the active site and catalyse the acyl-transfer reaction, a feature absent in the other members of the superfamily.

### A universal mechanism for rejection of highly abundant CoA in the alternative pocket

The primary attribute of a functional alternative pocket in FAALs is to discriminate and reject CoA from the chemically identical 4’-PPant of *holo*-ACP. Therefore, the pocket should allow the 4’-PPant entry into the tunnel but not the additional “head group”, adenosine 3’,5’-bisphosphate moiety of CoA. Structural analysis of 8 CoA-bound protein structures (constituting 14 protomers) of the ANL superfamily reveals the variability or degrees of freedom available for adenosine 3’,5’-bisphosphate moiety (head group). The analysis reveals that the 4’-PPant arm remains largely rigid in an extended form and the major source of variability is the conformational freedom around the 4’-phosphate (Supplementary Figure-6a). Such a variability has previously been used to classify the conformations of CoA as extended conformations (e.g.: *St*ACS, *Hs*FACL) or bent conformations (e.g.: *Aa*4’-CBL) ^31^. Limited variabilities in the conformations of the adenine ring and ribose are clustered in a small zone and therefore can be ignored (Supplementary Figure-6a). The length of the 4’-PPant arm is around 18-20 Å ^32^ while it is around 15-16 Å ^17^ as seen in the crystal structures of various ANL superfamily members, which is comparable to the length of predicted pockets (17-18 Å). The biochemical evidence (Figure-4c) along with length considerations indicates that the 4’-PPant arm can be accommodated within the predicted pocket. In the absence of structural information, the orientation of the 4’-PPant arm is random but limited to space with the identified pocket such that the 4’-phosphate is at the entry of the tunnel and the thiol near the active site.

Based on the above considerations, a CoA molecule in ANL superfamily members can be visualized as “flag hoisted on a mast”. The extended 4’-PPant arm can be considered as the “mast” and the adenosine 3’,5’-bisphosphate moiety as the “flag”. The rotation of the “flag” around the “mast” placed within the pocket is comparable to the degree of freedom available for the “flag” (Supplementary Figure-6b). We, therefore, rotated the “flag” around the “mast” to generate conformations at an angular sampling rate of 1° per conformation for each of the four known conformations of CoA (Figure-5a). It was found that main-chain atoms and Cβ atoms of the FAAL protein, irrespective of the conformation of the C-terminal domain (A-state or T-state), show an average of 28 clashes (van der Waals overlap > 0.25 Å) (Figure-5b). The adenosine 3’,5’-bisphosphate or the “flag”, would encounter severe clashes because the entry of the alternative pocket is deeply embedded within the subdomain-A and B of the N-terminal domain. It is unlike the entry of the canonical pocket, which is entirely open with the subdomain-B forming the base of the pocket (Figure-4). Therefore, the N-terminal domain of FAALs itself forms a strong deterrent for accommodating adenosine 3’,5’-bisphosphate and hence prevents CoA binding. Interestingly, the structural elements resisting the adenosine 3’,5’-bisphosphate are unique to FAALs such as the FSH (T252-R258 in *Ec*FAAL) and the loop harboring proline residues that guard the opening of the tunnel. Therefore, a strong negative selection and the absence of any positive selection make accommodation of adenosine 3’,5’-bisphosphate containing CoA unlikely. In contrast, the adenosine 3’,5’-bisphosphate lacking 4’-PPant of *holo*-ACP can easily access the pocket and participate in the acyl-transfer reaction. This essentially forms the structural basis for the CoA-rejection mechanism in FAALs (Figure-5c).

**Figure-5:**
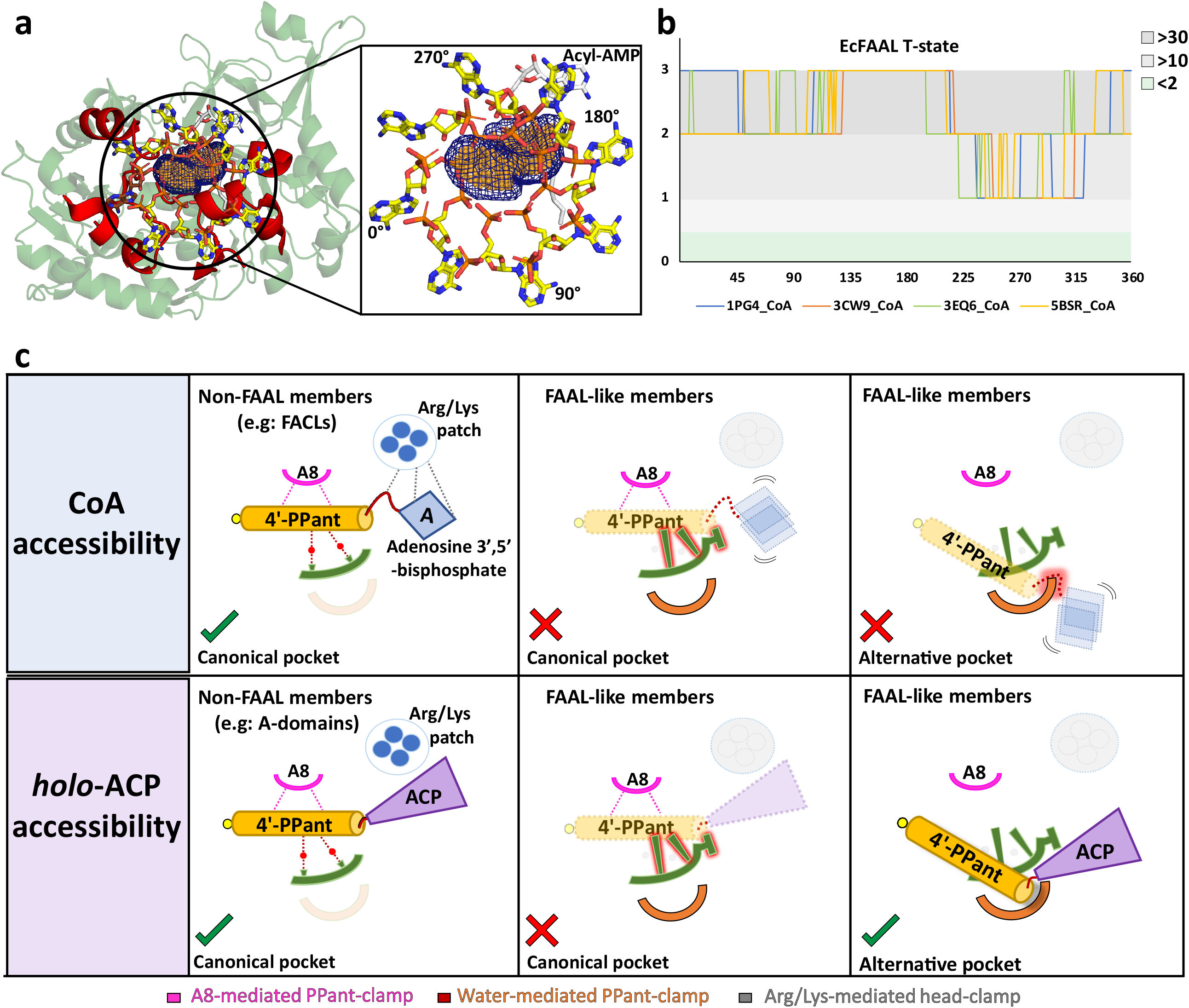
The alternative pocket in FAALs is highly selective, and its unique architecture negatively selects for CoA: (**a**) A “mast” (the 4’-PPant arm) is aligned along the longest length of the predicted pocket (blue; mesh representation) and the “flag” (adenosine 3’, 5’-bisphosphate) is rotated to generate theoretical orientations of the “flag”, These conformations of *Hs*FACL (PDB: 3EQ6) are shown (yellow; stick format) at 45° intervals. The regions of the *Ec*FAAL (cartoon representation) showing clashes are highlighted (red). (**b**) The count of atoms (C, N, O, Cα and Cβ) of *Ec*FAAL showing van der Waals short contacts (≤ 0.25 Å) with the conformations of the “flag” are scored (short contacts ≥ 30 = 3; ≥ 10 = 2; ≥ 2 = 1 and < 2 = 0). These scores (y-axis) are plotted against the specific rotation angles of the conformation, where the short contact is observed (x-axis). (**c**) The universal CoA-rejection mechanism is schematically summarised. The 4’-PPant (light orange) of both CoA and *holo*-ACP (purple) bind to the canonical pocket of non-FAAL members through multiple interactions (dotted lines). Interactions between Arg/Lys patch (circle; blue) and the adenosine 3’, 5’-bisphosphate moiety (rhombus; blue) acts as positive selection. The negative selection elements of the canonical pocket clash (highlighted in red) with the adenosine 3’, 5’-bisphosphate in the alternative pocket. The absence of Arg/Lys patch (dotted circle) fails to provide stability to the adenosine 3’, 5’-bisphosphate moiety (dotted rhombus; blue). The 4’-PPant (yellow) tethered to ACP (purple) is only accepted in the alternative pocket, which is absent in non-FAAL members, as it shows no clashes.

### Identification of FAAL-like proteins across different forms of life

Previous studies show that FAALs can be identified based on the presence of insertion and its anchorage to the N-terminal domain through hydrophobic interactions. A lack of sequence conservation in the FSI and only a single structural template for further analysis complicated the search procedure. Therefore, early genome mining efforts resulted in a sparse identification of FAALs in lineages of actinobacteria, cyanobacteria and proteobacteria along with some eukaryotes such as humans and mouse ^11^. The conserved sequence features from this study such as FSH, a blocked canonical pocket and a proline-lined 4’-PPant binding pocket led to the identification of FAAL-like proteins with greater confidence. The analysis revealed a ubiquitous distribution of FAALs across different forms of life including bacteria, plants, fungi, and animals, except archaea (Figure-6a). The phylogenetic analysis reveals that all the FAAL-like proteins diverge and cluster away from FACLs or A-domains. FAAL-like proteins show three sub-groups, viz. bacterial/plant FAALs group, fungal FAAL-like protein group and animal FAAL-like protein group (Figure-6a). Typical FAALs are bacterial FAALs, which are always found in a genomic context with PKS and PKS/NRPS hybrid genes either as a stand-alone domain or as a didomain fused to ACP or multidomain fused to the entire PKS/NRPS gene. Occasionally, the stand-alone bacterial FAALs are found to be interspersed with additional domains, mainly with dehydrogenases and oxygenases (Supplementary Figure-8b). Most plant FAAL-like domains resemble the bacterial FAALs from their sequence identity and domain organization. However, unique domain organizations are also found in plant FAAL-like domains, where an uncharacterized protein has FAAL-like domains sandwiched between HemY domains and catalase domains or amino acid oxidase domains (Supplementary Figure-8b). To the best of our knowledge, this is the first report of the presence of FAAL-like domains in plants and also the fusion of alternate domains to FAALs at their N-terminus.

**Figure-6:**
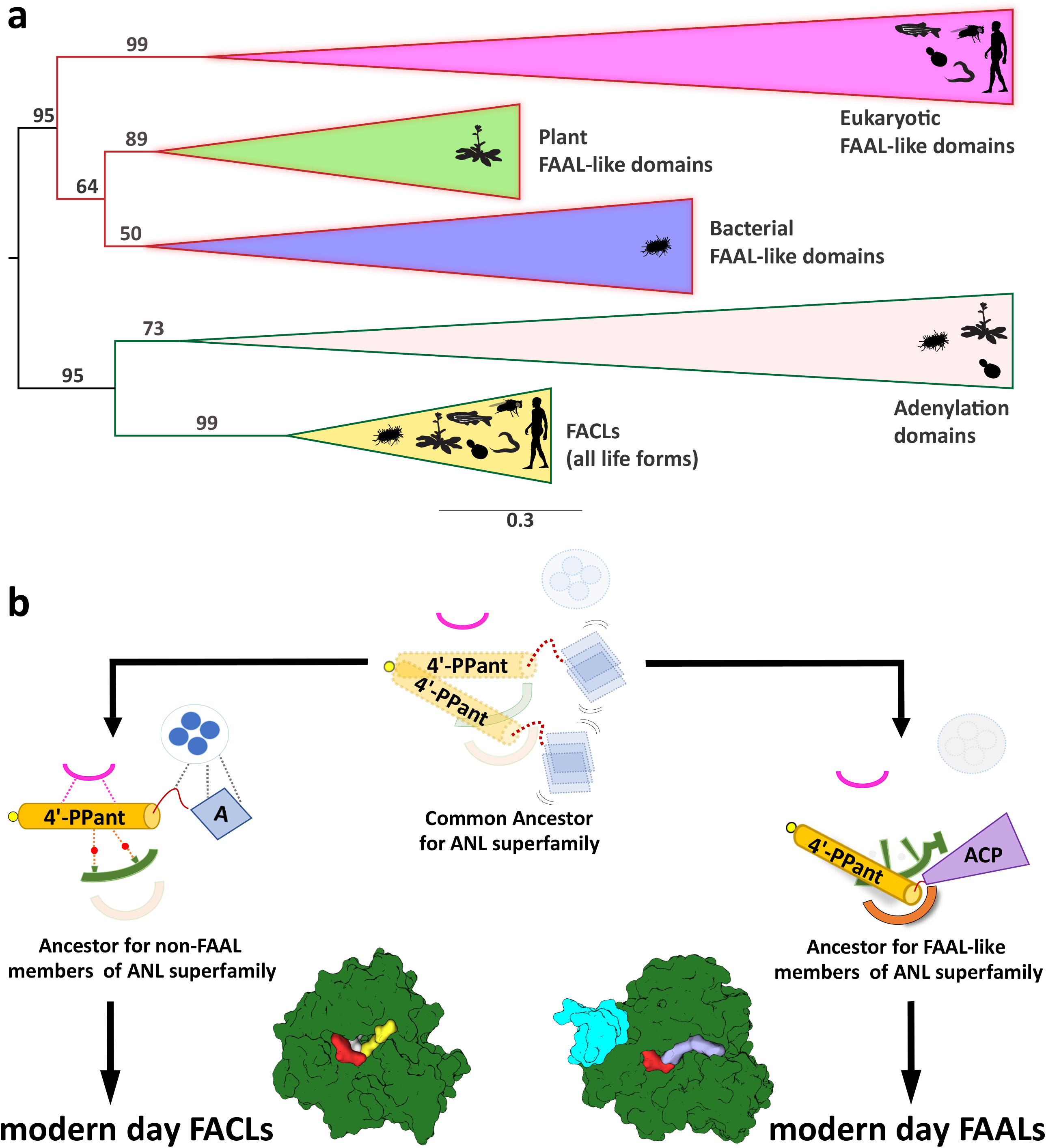
The CoA-rejection elements in FAALs are conserved in all forms of life and therefore, FAALs and FACLs have parallelly evolved from a common ancestor of the ANL superfamily: (**a**) A clustering diagram with the bootstrapping values for all the FAAL-like sequences is presented. The ANL superfamily members have two major divergent classes, viz., CoA-rejecting FAAL-like sequences and CoA-accepting non-FAAL members (FACLs and A-domains). A graphical representation of the distribution of these members is shown using pictorial representation of the organisms, which include bacteria, plants, yeast, worm, fly and human. (**b**) The ancestral fold of the ANL superfamily may have been a loose organisation of peptide scaffolds working with thiol-containing molecules such as pantetheine, etc. The ancestral fold accumulated various mutations resulting into the ancestor of modern-day acceptor-promiscuity lacking FAAL-like forms and acceptor-promiscuity containing non-FAAL members. The evolution of acceptor-promiscuity spectrum may have been driven based on their participation in bulk metabolic reactions. FAALs did not participate in bulk metabolic reactions and hence dedicated themselves to their cognate ACP partners to redirect small molecules to specific pathways. The non-FAAL members of the ANL superfamily such as FACLs participate in bulk reactions, where minor cross-reaction products are observed.

Fungal and animal FAAL-like domains tend to cluster together with bacterial/plant FAAL-like domains in bioinformatic analysis (Figure-6a) because of the conservation of unique FAAL-specific features. Despite their lower sequence identity, the FSI (Supplementary Figure-8b, 10c) supported by a hydrophobic patch and the FSH and CoA-rejecting features (Supplementary Figure-9a, 11b) are clearly identifiable. Such conserved FAAL-specific features indicate that FAAL-like modules are recruited in eukaryotes for specific metabolic processes owing to their unique ability to reject CoA. Interestingly, fungal and animal FAAL-like domains share a highly conserved three-domain architecture, and such an architecture is not seen in any of the prokaryotes and plants (Supplementary table-III). The conserved three-domain protein consists of a N-terminal DMAP1-binding domain followed by a tandemly fused two FAAL-like domains as found in Fungi to Animals. The eukaryotic metabolic context for avoiding a reaction with CoA through these FAAL-like domains and their unique domain organization remains to be explored. Based on these observations, FAAL-like domains are more wide-spread than previously anticipated and their omnipresence is comparable to the universally present FACLs. Therefore, we propose that FAALs may not have descended from FACLs and they rather share a parallel evolutionary history. The ancestral ANL fold could have diverged as non-promiscuous FAAL-like members and promiscuous nonFAAL-like members simultaneously in the last universal common ancestor (Figure-6b).

## CONCLUSIONS

The study uncovers the mechanistic basis of how FAALs strictly reject CoA, which is highly abundant and almost chemically identical to their actual substrate, *holo*-ACP. The rejection mechanism relies on a discriminatory 4’-PPant-accepting pocket while avoiding the promiscuous canonical CoA-binding pocket that is rendered non-functional. It has been achieved through bulky hydrophobic residues in the pocket and a unique secondary structural element, FSH at the entrance. The discriminatory 4’-PPant accepting pocket in FAALs, on the other hand, has a unique architecture that negatively selects adenosine 3’, 5’-bisphosphate moiety and also lacks Arg/Lys residues for positive selection.

Interestingly, these rejection criteria are not only conserved in bacteria but also in all forms of life (excluding archaea). The unique remodelling of pockets to ensure acceptor discrimination probably allowed the evolutionary recruitment of FAAL-like domains in metabolic crossroads for redirecting the fate of molecules to specific pathways. Such a wide-spread conservation of FAAL-like proteins puts the evolutionary origin of FAALs parallel with FACLs in the last universal common ancestor and not as a subset of FACLs.

The scaffold of the ANL superfamily of enzymes is known to be promiscuous not only for the substrates they act on ^33^ but also the final acceptor of the acyl adenylates. Surprisingly, billions of years of evolution has not prevailed upon the substrate-promiscuity problem as well as the acceptor-promiscuity problem. The probability of acceptor-promiscuity influencing the erroneous product formation is high as some of the acceptors such as CoA and pantetheine are the most abundant molecules in the cell. The problem is greatly amplified as it can redirect the fate of metabolites from one pathway to another such as primary metabolism to secondary metabolism. Our analysis shows that none of the solved CoA-bound structures of ANL superfamily members, even the protomers of the same crystal (e.g., *Se*FACL), exhibit conformity in the binding mode of 4’-PPant or the adenosine 3’,5’-bisphosphate moiety ^15,34^. The variability in the 4’-PPant binding is also true for the A-domain:ACP complexes ^17,35,36^. Hence, it is evident that a defined CoA-binding pocket is lacking in ANL superfamily members, which is commensurate with the failure to identify the pocket in FACLs using various pocket-search algorithms. Therefore, arbitrary access of the active site without any selection determinants is likely to be the root cause of final acceptor promiscuity in the case of these enzymes.

It is not clear if these observed spectrum of latent activities in these enzymes’ design is an evolutionary relic or has any physiological relevance in a specific cellular context. The persistence of acceptor promiscuity can only have two explanations: viz., the cross-reaction products are beneficial, or pocket modification is not possible without compromising the basic function. In this context, FAALs are surprisingly high-fidelity enzymes representing the extreme end of the promiscuity spectrum, offering no cross-reaction with CoA as acceptors of the acyl adenylates ^9-11,37^. A functional and discriminatory 4’-PPant binding in FAALs have established them as an alternative enzymatic bridge between FA synthesis, exogenous fatty acids import and PKS/NRPS machinery. FAALs act as a loading module in PKS/NRPS-mediated biosynthesis of diverse bioactive natural products with fatty acids such as mycosubtilin, daptomycin, micacocidin, ralsomycin, olefin, tambjamine, ambruticin, puwainphycins, jamaicamide, columbamides to name a few. The relevance of the lack of acceptor promiscuity in FAALs has been demonstrated in Mycobacteria as being responsible for dictating the fate of free-fatty acids in producing virulent lipids ^10^.

The conserved sequence and structural features of bacterial FAALs enabled better identification of FAAL-like domains across all forms of life, except archaea. FAALs may have been recruited to specifically “load” fatty acids on PKS/NRPS but the possibility of lack of promiscuity in FAALs driving their over representation cannot be ignored. Their characteristic absence in archaea perhaps can be explained by the abysmal frequency or absence of PKS or NRPS systems. FAAL-like domains of bacteria and plants exhibit remarkable sequence similarity, which is expected because of the presence of canonical PKS/NRPS enzymology in both systems. A few of the identified FAAL-like domains in bacteria show unique domain organizations such as a fusion with lysophospholipid acyltransferases and alpha aminoadipate reductases or found interspersed in operons with dehydrogenases, decarboxylase and ketoacyl synthases. Unique domain architectures are also found in plants, where FAAL-like domains are sandwiched between HemY and catalase domains and the function of these conserved proteins remain unknown. These unique architectures are evolutionary instances of the recruitment of FAAL-like domains for PKS/NRPS independent functions.

Interestingly, unique domain architectures are found in FAAL-like domain-containing proteins of fungi and animals, the majority of which are largely PKS/NRPS free systems. In these organisms, two tandemly fused FAAL-like domains are found at the C-terminus and a DMAP-1 binding domain at the N-terminus of a highly conserved protein. These proteins are annotated and characterized as the virulent factor CPS1 in few fungal pathogens such as *Magnaporthe oryzae* ^38^, *Cochliobolus heterostrophus* ^39^ and *Coccidioides posadasii* ^40^. Therefore, they are proposed as a target for potential fungicides and as a vaccine candidate in plant and animal pathogens respectively ^38-40^. In invertebrates and vertebrates, these are known as Dip2 and are important for proper axon bifurcation and guidance suggesting their physiological importance ^41,42^. Recently, we showed that the mice lacking one of the eukaryotic FAAL-like proteins, Dip2a, show a diet-dependent growth anomaly such as obesity pointing to their importance in eukaryotic lipid metabolism ^43^. The precise molecular and biochemical details of these proteins are yet to be elucidated. Given the extent of sequence divergence in various motifs of eukaryotic FAAL-like domains, it is difficult to predict the processes they may be involved in. The presence of several elements of the CoA-rejection mechanism and their close identity to FAALs allow us to put forth the hypothesis that eukaryotic FAAL-like domains were recruited for their high acceptor fidelity property. It is also possible that these divergences have some functional relevance in the context of eukaryotic metabolism, which may represent an additional member in the spectrum of biochemical activities represented in the ANL superfamily.

The current work identifying FAAL-like enzymology using a universal CoA rejection mechanism may form the platform for further studies to delineate why they have been recruited in fungal and animal systems. The study provides new structural and sequence attributes to confirm the identity of FAALs, many of which remains misannotated and uncharacterized. The study opens new avenues in combinatorial engineering of PKS/NRPS by using FAALs as a unique module to load fatty acids or engineer them to load unique molecules with exceptional fidelity. Thus, FAALs can be exploited to produce novel bioactive molecules by virtue of their unique acceptor-fidelity property.

## MATERIAL AND METHODS

### Cloning, expression and purification of proteins

*Ec*FAAL (A0A0H2VDD9), *Ec*ACP (A0A2×1NC35), *Ec*FACL (P69451), *Ms*FAAL32 (A0R618), *Ms*PKS13 (A0R617; 1-1042 residues), RsFAAL (Q8XRP4), *Rs*ACP (Q8XRP0), *Af*FACL (O30147), *Mx*FAAL (Q1CXX0), *Mx*ACP (Q1CXW9) and *Mt*FACL13 (P9WQ37) were amplified by PCR from their respective genomic DNA using Phusion polymerase (Thermo Scientific). These genes were expressed as hexahistidine-tagged proteins after the induction with IPTG using the *E. coli* BL21(DE3) expression system. The mutants were generated using quick-change site-directed mutagenesis. All the proteins including mutants were expressed and purified to homogeneity using Ni-NTA affinity chromatography followed by size-exclusion chromatography at 4°C. They were flash-frozen in liquid nitrogen and stored at -80°C until further use.

### Biochemical analysis of FAALs and FACLs

The acyl-AMP and acyl-CoA formation by FAALs and FACLs were performed by previously described methods ^10^. All FACLs (*Mt*FACL13, *Ec*FACL, *Af*FACL), EcFAAL and MsFAAL32 were used at 5 μM concentration while MxFAAL and RsFAAL were used at 7.5 μM. The ^14^C-fatty acids allowed detection of the products on a phosphorimager (Amersham Typhoon FLA 9000), which were then quantified by densitometry using Image Lab (Bio-Rad Laboratories Inc.). All experiments were performed as triplicates. The percentage of acyl-AMP converted to acyl-CoA from the total acyl-AMP formed is used to plot and compare the activity, along with the standard error of the mean, of wild-type against the respective mutants.

### Conversion of *apo*-ACP to *holo*-ACP

All the purified ACPs were converted to *holo-*ACP before being flash-frozen using previously described protocols ^44^. Briefly, after Ni-NTA purification, they were buffer exchanged to the phosphopantetheinylation buffer (20 mM Tris pH 8.8, 10 mM MgCl_2_ and 10 mM dithiothreitol). ∼250 μM of ACP was then incubated with 1.25 μM of a non-specific phosphopantetheinyl transferase Sfp from *B. subtilis* in a 300 μl reaction mixture along with 10 mM MgCl_2_ and 1 mM CoASH. The reaction mixture was incubated at 25°C for 12-16 hours and further purified using size-exclusion chromatography at 4°C. The *holo*-ACP was flash-frozen in liquid nitrogen and stored at -80°C until further use. The conserved serine on which the 4’-PPant moiety is added, post-translationally, is mutated to alanine and the resulting protein is used as *apo*-ACP for all assays.

### Loading activated acyl chains on *holo*-ACP by FAALs

The transfer of activated acyl-chain to stand-alone *holo*-ACP by FAALs was assessed using radiolabelled fatty acids combined with conformationally-sensitive Urea-PAGE ^30^. The following ratio of FAAL to *holo*-ACP was used: 1 μM of *Ec*FAAL with 20 μM of EcACP, 4 μM of *Mx*FAAL with 8 μM of MxACP, 5 μM of RsFAAL with 20 μM RsACP and 1 μM of *Ms*FAAL32 with 12.5 μM of *Ms*PKS13ΔC was used. These proteins were incubated in a typical 15 μl reaction mixture composed of 50 mM HEPES pH 7.2, 5 mM MgCl2, 0.01% w/v Tween-20, 0.003 % DMSO, 2 mM ATP and 9 μM of ^14^C labelled-fatty acid. The reactions of *Mx*FAAL-MxACP and *Rs*FAAL-*Rs*ACP consisted of 20 mM Tris pH 8.0 and lacked DMSO. The reaction mixture was incubated for 2 hours at 30°C and quenched with an equal volume of urea loading dye (25 mM Tris pH 6.8, 1.25 M urea, 20% glycerol and 0.08% Bromophenol blue). The contents were immediately loaded and separated on a 15% PAGE containing 2.5 M urea. The acyl-transfer on *holo*-*Ms*PKS13_1-1042_ by *Ms*FAAL32 was carried out as previously described ^29^. Briefly, ^14^C labelled-fatty acid was used to assess the transfer on *holo*-*Ms*PKS13_1-1042_ on an 8% SDS-PAGE. All gels were then dried using a gel drier (Bio-Rad gel dryer 583) and the radiolabelled *acyl*-ACP was detected by using the phosphorimager (Amersham Typhoon FLA 9000).

### Sequence and structural analysis

All sequences were identified and retrieved from the NCBI sequence database using EcFAAL as template and BLAST search algorithm. A structure-based sequence alignment was generated using msTALI ^45^ and then other sequences were added to the alignment in MAFFT ^46^. The sequence alignments were rendered using ESPript ^47^. The phylogenetic analysis of these sequences was carried out using both neighbour-joining as implemented in MAFFT^46^ and maximum-likelihood as implemented in IQTREE ^48, 49^ using default parameters. The phylogenetic tree was rendered in MixtureTree Annotator ^50^. The images of various organisms were reusable silhouette images of organisms obtained from Phylopic (www.phylopic.org) under Creative Commons license. All structural analysis including structural superposition, van der Waal distance measurements, ligand alignment was carried out with various in-built features of PyMOL ^51^. All cavity search programs were run using default parameters.

## AUTHOR CONTRIBUTIONS

GSP and PK contributed equally to the work by designing and performing all the experiments. SP, ND, SM and SS also contributed to expression and purification of protein. SN contributed to initial optimization of acyl-transfer assays of FAALs. MKM, KDP, SM and SS contributed to generation of constructs. RSN conceived and supervised the study. GSP, PK and RSN wrote the manuscript with help from RSG, BP and SM. All the authors have analysed and reviewed the data and manuscript.

## FUNDING SOURCES

GSP and PK thank the Department of Biotechnology, India for research fellowship. SM and SS thank CSIR, India and UGC, India for research fellowship. R.S. thanks SERB-NPDF, NCP under health care theme project of CSIR, India; J.C. Bose Fellowship of SERB, India; and Centre of Excellence Project of Department of Biotechnology, India.

## ACKNOWLEDGMENT

We acknowledge Dr. Pananghat Gayathri (IISER-Pune) and Dr. Raghunand R. Tirumalai (CSIR-CCMB) for sharing the genomic DNA samples of *Myxococcus xanthus* and *Mycobacterium smegmatis*, respectively.

## COMPETING INTERESTS

The authors declare that they have no competing interests.

## DATA AND MATERIALS AVAILABILITY

All data needed to evaluate the conclusions in the paper are present in the paper and/or the Supplementary Materials. Additional data related to this paper may be requested from the authors.

## Notes

### Competing Interest Statement

The authors have declared no competing interest.

