## supplementary figure 1-10 for "A universal pocket in Fatty acyl-AMP ligases ensures redirection of fatty acid pool away from Coenzyme A-based activation"

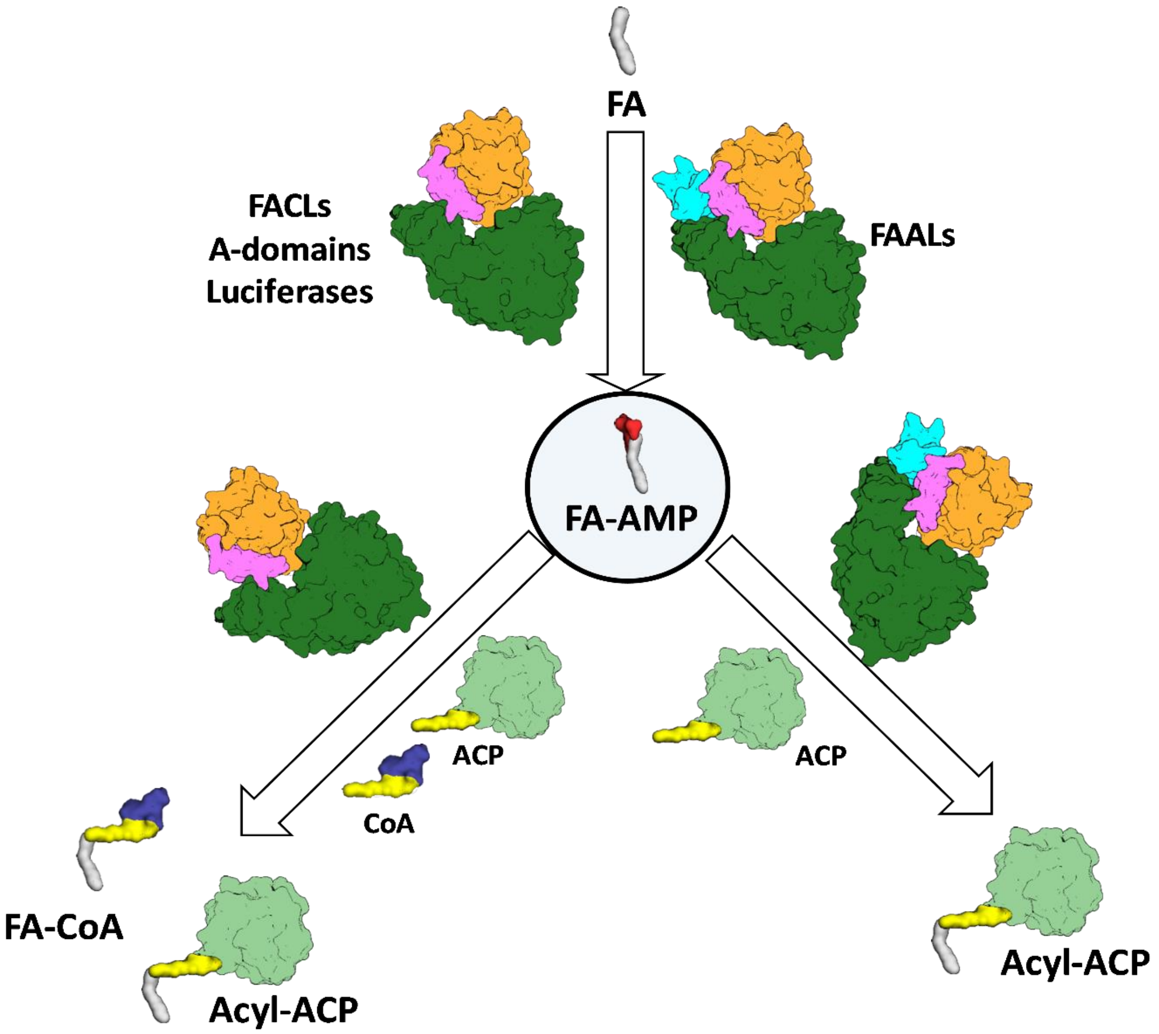


**Supplementary Figure-1**: An overview of the characteristic enzymology presented by FAALs and FACLs leading to the dichotomy of the fate of free-fatty acids. FAALs and FACLs, both produce the fatty acyl-AMP from fatty acid (grey) but transfer the intermediate to different acceptors. Such a differential activity is despite two critical similarities; firstly, the overall structural similarity (N-terminal domain in green; C-terminal domain in orange connected by the A8-motif in violet), except an insertion in FAALs (cyan). Secondly, the differential activity is despite the chemical identity of the substrates that react during the second step, where CoA used by FACLs is used to post-translationally modify *apo*-ACP to *holo*-ACP (light green) that utilized by FAALs. FAALs are highly specific in diverting the intermediate fatty acyl-adenylate to the *holo*-ACP acceptor and then divert the fate of the pool of free-fatty acids towards various pathways that work independently of CoA.

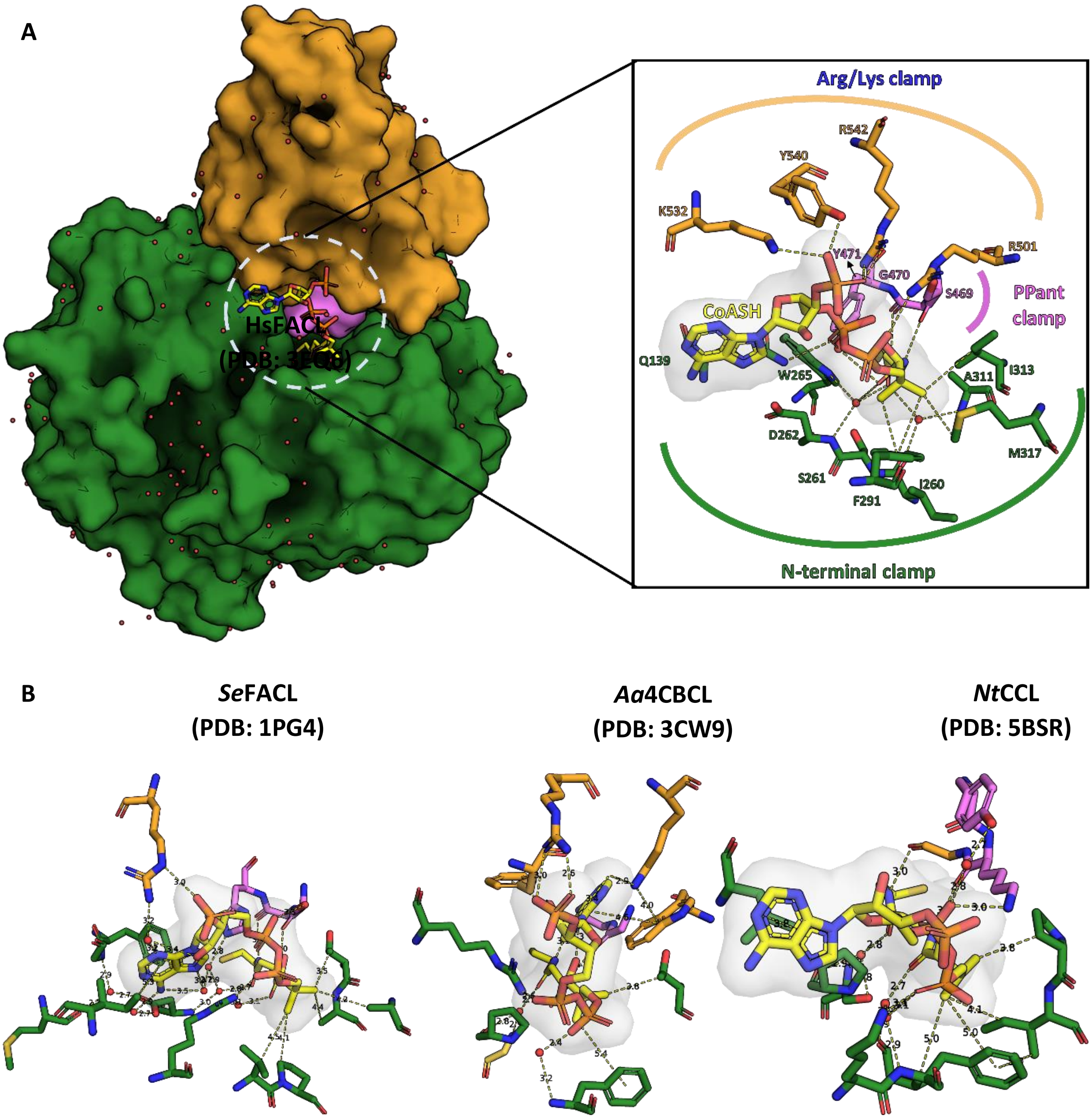


**Supplementary Figure-2: (a)** A surface representation of CoA-bound *Hs*FACL (PDB: 3EQ6) is shown with CoA (yellow) as a stick model sandwiched between the N-terminal (green) and C-terminal domain (orange) along with the connecting A8-motif (violet). The inset highlights the network of interactions between the CoA and the protein molecule, which can be grouped into three categories. (i) Arg/Lys clamp: Structurally conserved Arg/Lys residues from the C-terminal domain (occasionally from the N-terminal domain) make ionic interactions with phosphates of CoA. The sequence positions of these positively charged residues need not be conserved. (ii) PPant clamp: Residues from the highly conserved A8-motif form hydrogen bonds with the 4'-phosphopantetheine arm. (iii) N-terminal clamp:

A series of water-mediated interactions with residues from the N-terminal and CoA. **(b)** A stick model of the CoA and residues of the protein forming a network of interactions is shown from three diverse CoA-bound FACL structures, namely acetyl-CoA synthetase (*Se*FACL; PDB: 1PG4), 4-Chloro benzoyl-CoA ligase (*Aa*FACL; PDB: 3CW9) and Coumarate-CoA ligase (*Nt*CCL; PDB: 5BSR). The residues from the C-terminal domain are shown in orange, those from the N-terminal domain are shown in green and residues from A8-motif are shown in violet. These interactions highlight the plasticity of the CoA-binding pocket with limited or no specific residues that impart specificity for CoA.

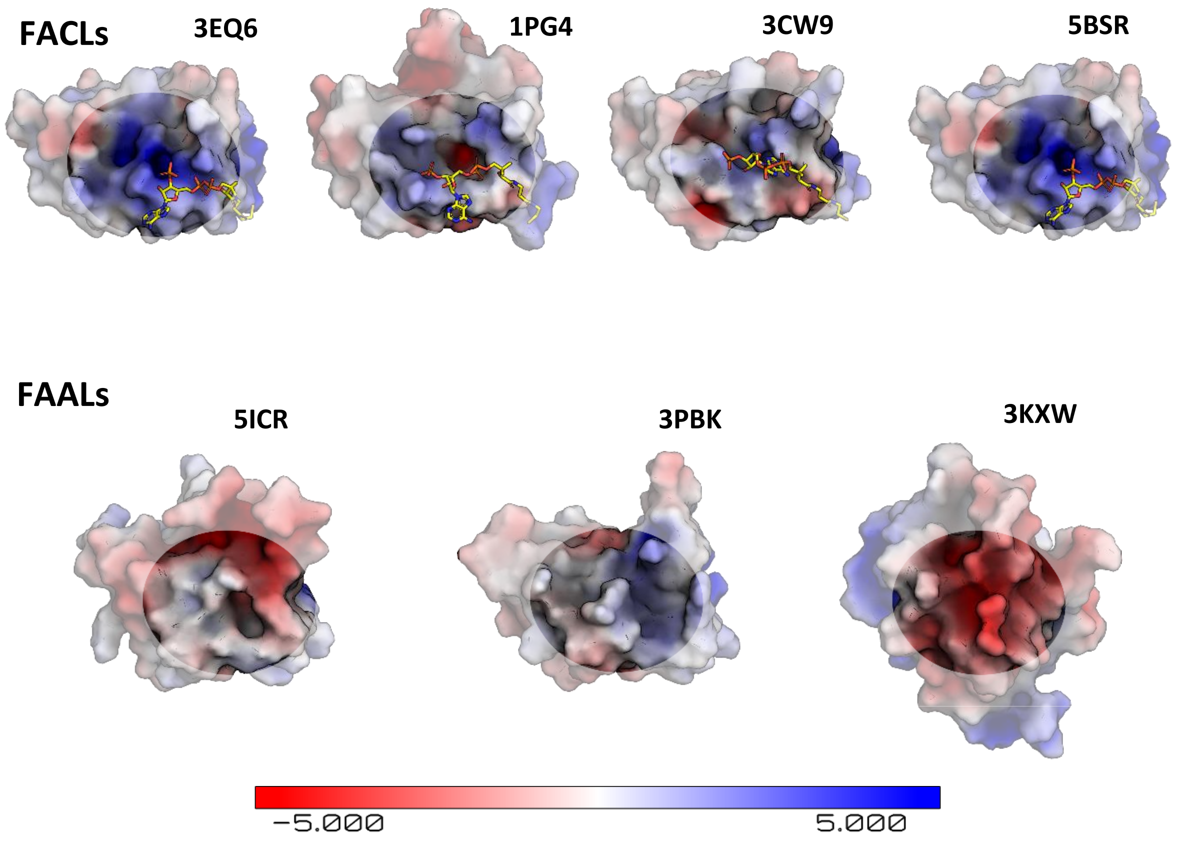


**Supplementary Figure-3: (a)** An electrostatic map of the C-terminal domain from CoA-bound FACLs structures is shown along with a stick model for CoA, which shows that there are positive rich regions that assist in holding the phosphates of CoA. **(b)** The electrostatic potential map of the crystal structures of the C-terminal domain of FAALs is shown for comparison, where the structurally analogous region binding to CoA is devoid of positively charged patch as seen in the case of FACLs. The electrostatic map was generated using the APBS module in PyMOL[51].

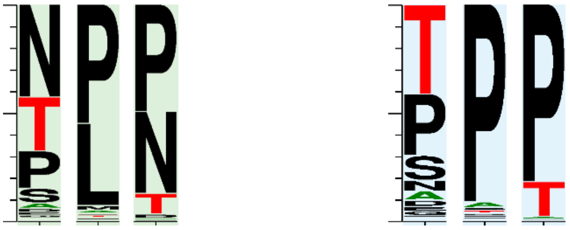

**Supplementary Figure-4:** A comparison of the frequency of the prolines surrounding the newly identified alternative pocket in FAALs (highlighted in blue) against the frequency of prolines in FACLs (highlighted in green). The representation indicates that the regions are highly variable. The conservation of prolines is higher in FAALs than FACLs. It may be occasionally replaced with Threonine or Serine in case of FAALs and in case of FACLs it is replaced by Leucine or Asparagine. The frequency plot was generated from the sequence alignment using web-LOGO[1].

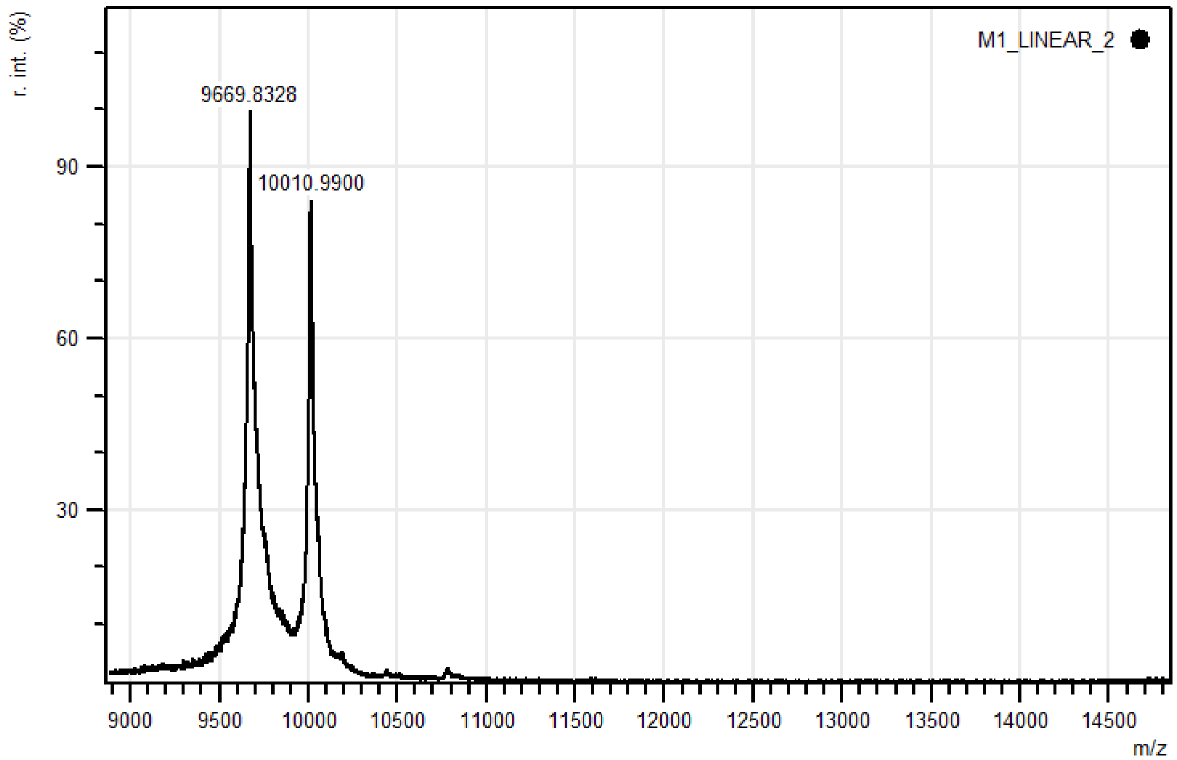
r. int. (%)
M1_LINEAR_2
9669.8328
10010.9900
90
60
30
9000
9500
10000
10500
11000
11500
12000
12500
13000
13500
14000
14500
m/z

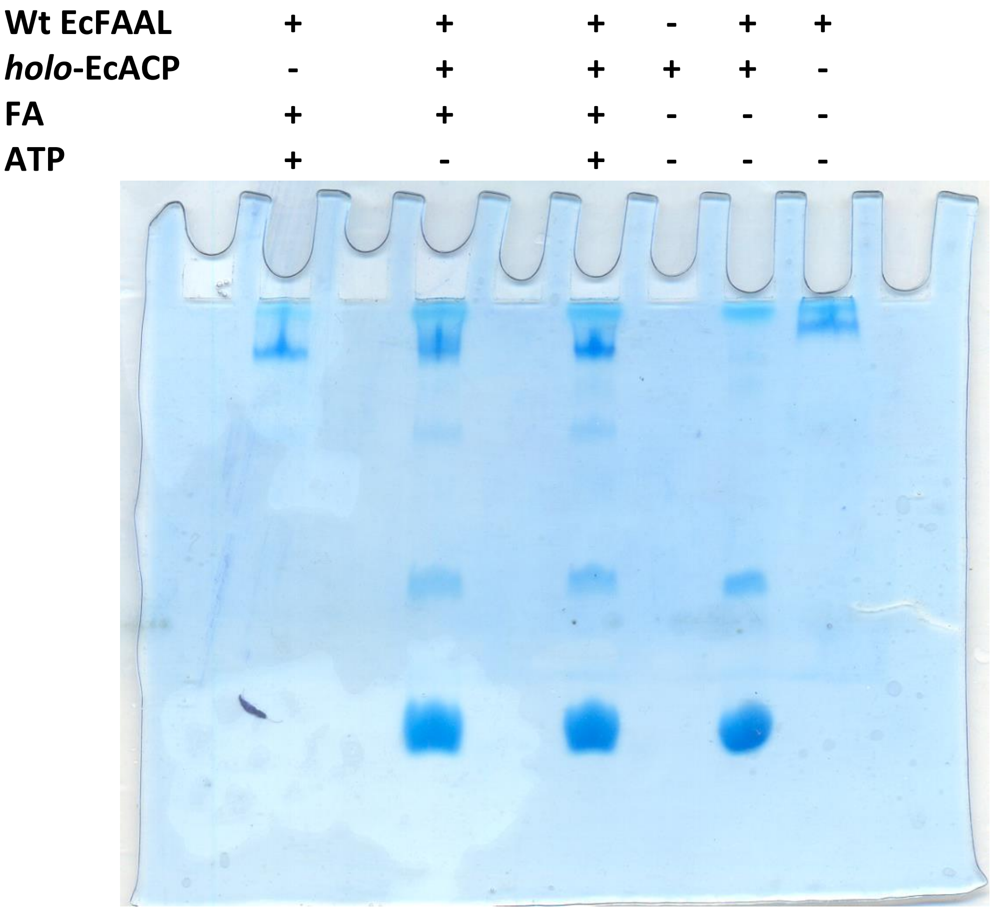
Wt EcFAAL + + + - + +
holo-EcACP - + + + + -
FA + + + - - -
ATP + - + - - -

**Supplementary Figure-5:** Typically, acyl-transfer reactions with ACP are assessed using an SDS-PAGE (coupled to radiolabelled or fluorescently labelled substrates) or conformationally-sensitive urea-PAGE or HPLC-coupled to mass spectrometry. SDS-PAGE based assays are suitable for ACPs that are fused to PKS/NRPS modules, which was used for MsFAAL32-MsPKS13$_{1\text{-}1042}$ in this study. This strategy is not suitable for smaller-sized molecules such as ACP (<10 kDa), which often migrate at the edges along with residual labelled fatty acid and acyl-adenylates. The assay also requires converting *apo*-ACP to *holo*-ACP using an enzyme, Sfp from *Bacillus subtilis*. **(a)** A representative image of the mass-spectrometry analysis (AB-SCIEX 4800 MALDI-TOF and using 5 mg/ml sinapinic acid in 50% acetonitrile and 0.1% trifluoroacetic acid as matrix) of *Sfp* catalysed post-translational modification of EcACP (9.6 kDa) to form *holo*-ACP (10 kDa). The analysis revealed that less than 50% of the *apo*-ACP is only converted to *holo*-ACP. **(b)** A representative image of Coomassie-stained conformationally sensitive Urea-PAGE (15% Acrylamide and 2.5M Urea) of the EcFAAL catalysed acyl-transfer on *holo*-EcACP. The Coomassie-stained gel does not reveal the classic separation of *holo*-ACP and *acyl*-ACP.

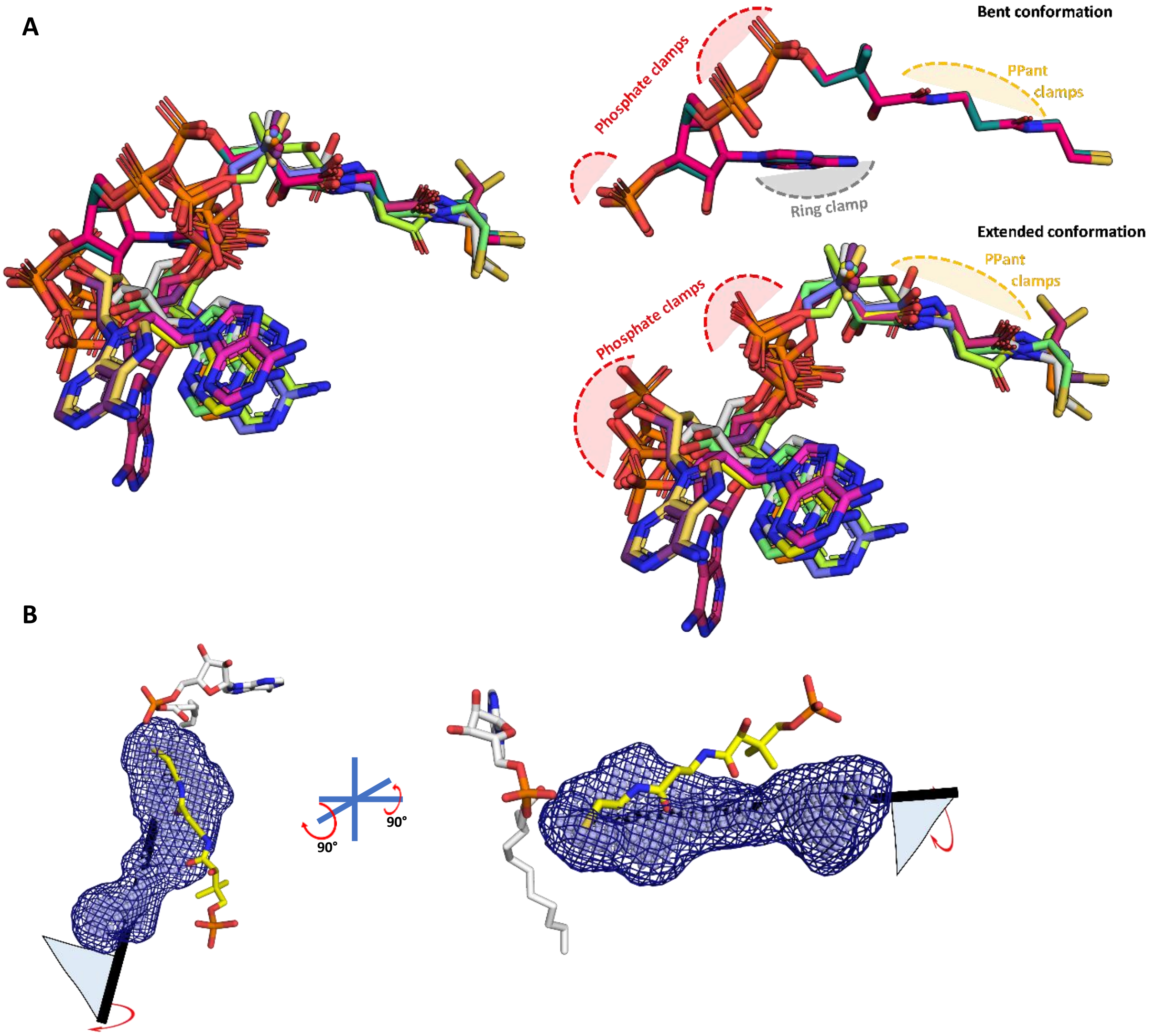


**Supplementary Figure-6: (a)** The available CoA-bound crystal structures of the ANL superfamily members were taken, which include *St*ACS (1PG4, 2 protomers; 2P2F, 2 protomers; 2P2J, 2 protomers), HsACSM2A (3EQ6, 2 protomers), *Nt*CCL (5BSR, 1 protomer), *Ds*PrpE (5GXD, 1 protomer) and *Aa*4CBL (3CW9, 2 protomers). These were aligned based on the maximum-common-substructure method as implemented in *mcsalign* plugin of PyMOL[51] along the length of the 4'-PPant. The superposition reveals that CoA structures are bound to ANL superfamily members mainly in an extended conformation (adenine ring away from the 4'-PPant arm) and rarely in bent conformation (adenine ring in proximity to 4'-PPant arm). The difference is due to the conformational freedom around the phosphodiester bonds, however, despite the variations, the phosphates tend to spatially organize in the similar positions available for the phosphate clamping Arginine or Lysine residues. The bent conformation has an additional contact point, ring clamp, where the adenine ring is held through aromatic amino acids, while the adenine ring is free in the extended conformation of CoA with orientations of adenine ring having a degree of variability but restricted to a small zone. The 4'-PPant arm is presented in a fully extended form in both the subsets of CoA and the PPant clamp atoms presented for hydrogen bonding with the A8 motif and water molecules. (**b**) In the absence of any structural information on how the 4'-PPant may be accommodated in the pocket, it is safe to assume the 4'-PPant is accommodated within the identified pocket analogous to the "mast" of the flag. The flag here represents the head group, which can be rotated to identify any possible orientation that can be accommodated within the alternative pocket. All the known conformations of head groups from CoA-bound ANL superfamily members serve as a template in this case.

A

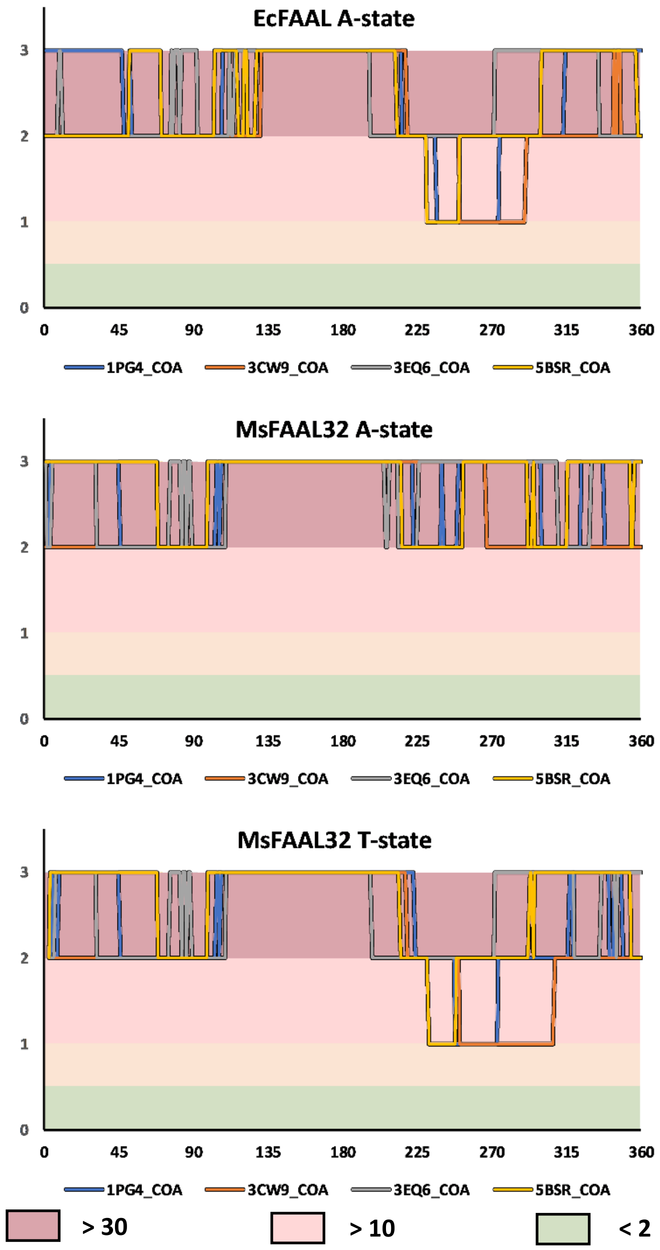
EcFAAL A-state
MsFAAL32 A-state
MsFAAL32 T-state
0
1
2
3
0
45
90
135
180
225
270
315
360
1PG4_COA
3CW9_COA
3EQ6_COA
5BSR_COA
> 30
> 10
< 2

B

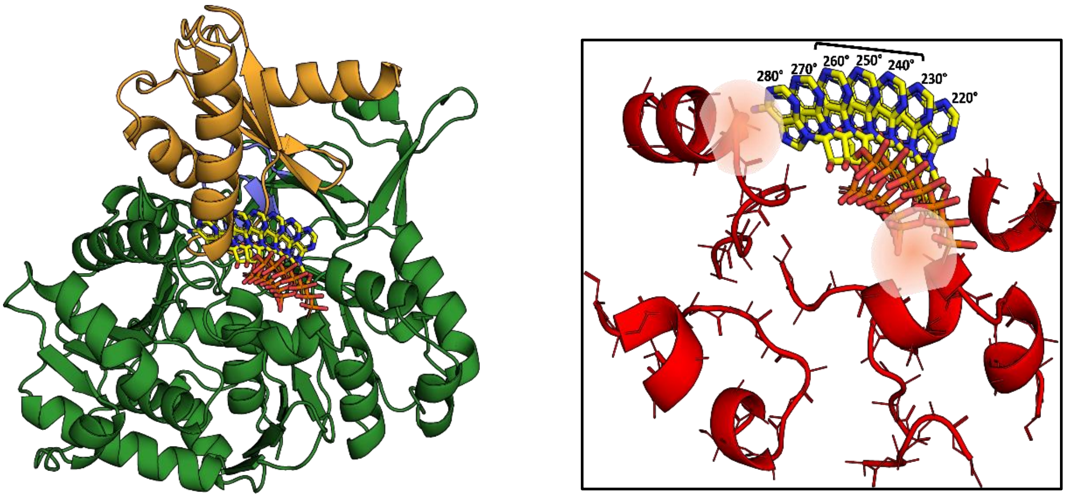
280°
270°
260°
250°
240°
230°
220°

**Supplementary Figure-7: (A)** The variation of clash score with an angle for the possible adenylation and putative thioesterification conformations of the two known FAALs structures (*Ec*FAAL; PDB:3PBK and MsFAAL32; PDB: 5ICR) is presented. It is very clear that none of the conformations, A-state or T-state, of both the proteins, *Ec*FAAL and *Ms*FAAL32, are capable of accommodating any of the known conformations of adenosine-3',5'-bisphosphate moiety of CoA. A large number of the sampled conformations present clashes with more than 30 mainchain atoms even with a conservative distance of ≤ 0.25 Å. The plot for *Ec*FAAL in A-state and T-state is comparable in the zones of least resistance, which is also true for *Ms*FAAL32 T-state suggesting that these are conformation-independent. This observation indicates that the clashes can primarily be attributed to the N-terminal itself and the unique location of the pocket. It should be noted that the plot for *Ms*FAAL32 A-state is unique because of the alternative pocket is occluded by a FAAL32-specific insertion described by Aldrich C.C., Anderson W.F and co-workers. **(B)** The T-state *Ec*FAAL was modelled by superposing the C-terminal domain with HsACSM2A and show in cartoon form (N-terminal in green and C-terminal in orange) along with theoretical orientations of the head group CoA from 220°-280° is shown in stick representation (yellow). The major elements of the N-terminal domain in *Ec*FAAL that contribute to clashes are shown in a cartoon representation (red) along with the stick representation of the mainchain atoms and Cβ. The conformations of the head group from 220°-280°, where only a few (2-10) clashes are observed, particularly the region spanned between, 240°-260°. The zone of least resistance is largely owing to a vacant space between N- and C-terminal domains without any residues to hold the phosphates or adenine ring. The space occupied by these conformations is close to the typical structural site for CoA accommodation in FACLs except that the adenine ring is oriented away from the protein. The typical Arg/Lys residues that bind phosphates are absent in FAALs and hence these conformations cannot be expected to be stabilized by phosphate clamps. The orientations with inward-facing adenine ring resemble the bent conformation of CoA, a rare conformation in ANL superfamily members, where the adenine ring is held by a conserved tryptophan as seen in *Aa*CBL (PDB: 3CW9). The conserved tryptophan residue is absent in most ANL superfamily members including FAALs and FACLs and hence these conformations cannot be expected to be stabilized by stacking interactions with adenine. Therefore, in the 220°-280° region even though the rejection by negative selection is weak, there is no scope of positive selection which renders the region incompatible with CoA-binding.

**A**

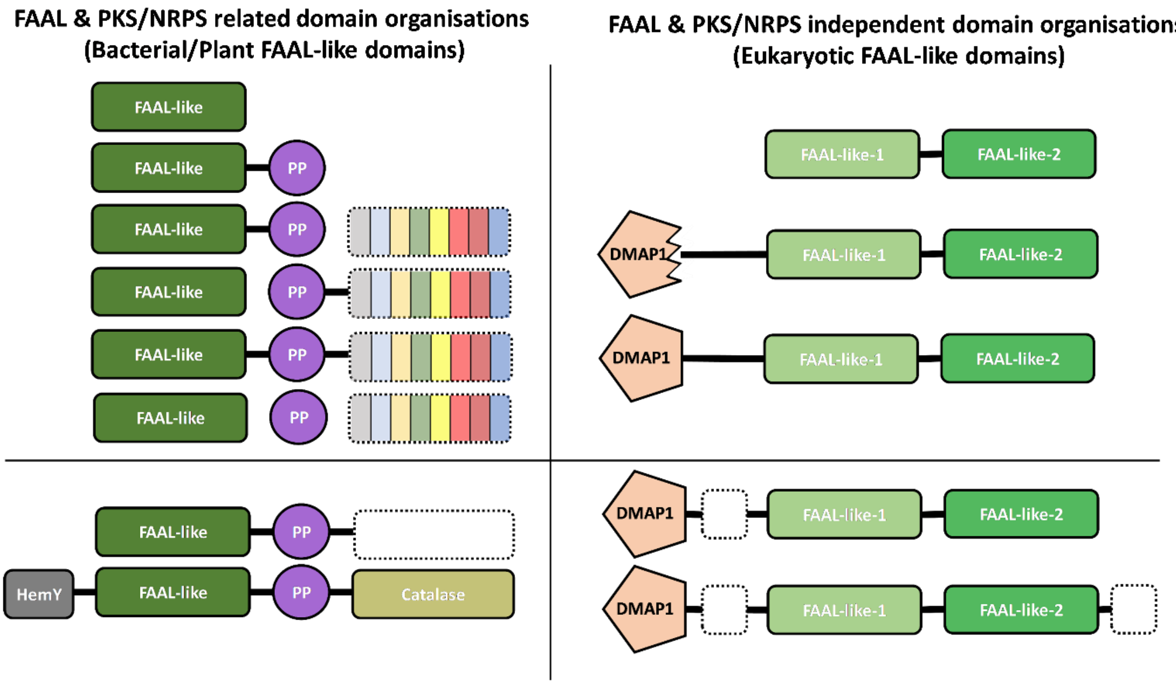

FAAL & PKS/NRPS related domain organisations
(Bacterial/Plant FAAL-like domains)
FAAL & PKS/NRPS independent domain organisations
(Eukaryotic FAAL-like domains)
FAAL-like
PP
HemY
Catalase
DMAP1
FAAL-like-1
FAAL-like-2

**B**

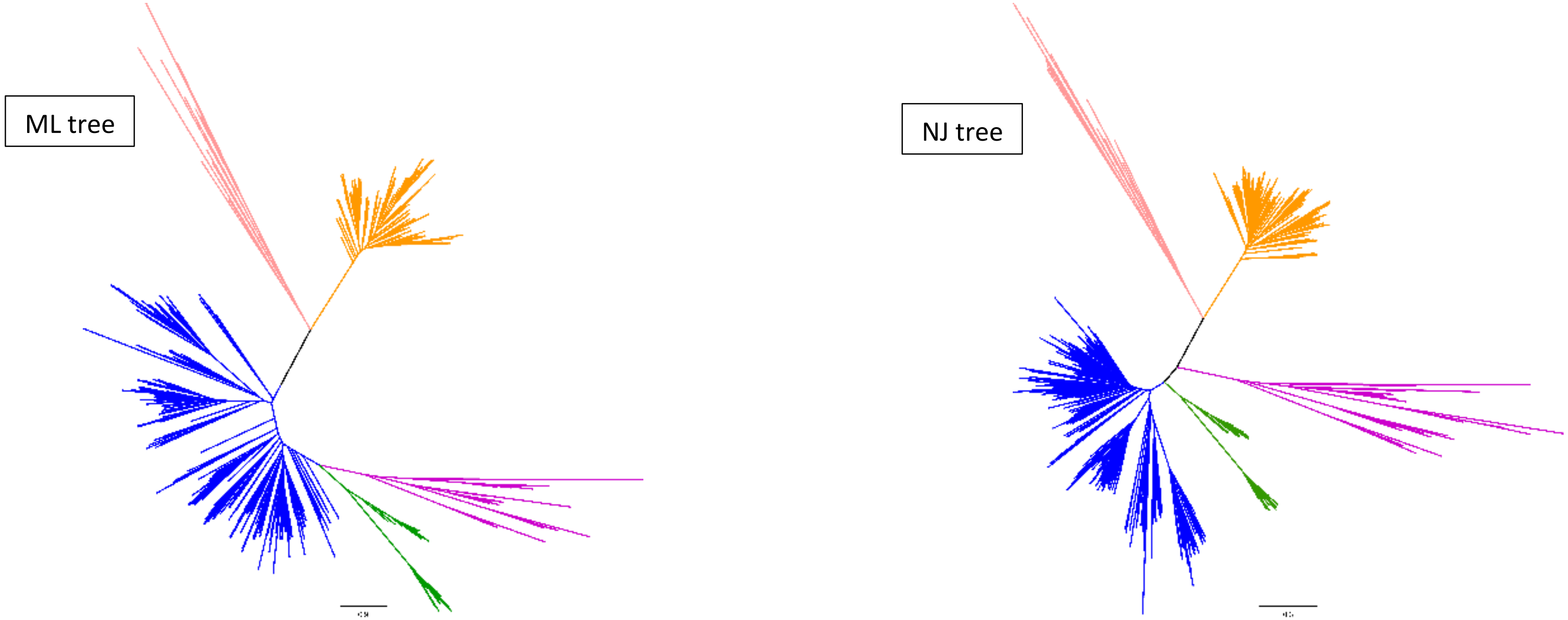

C

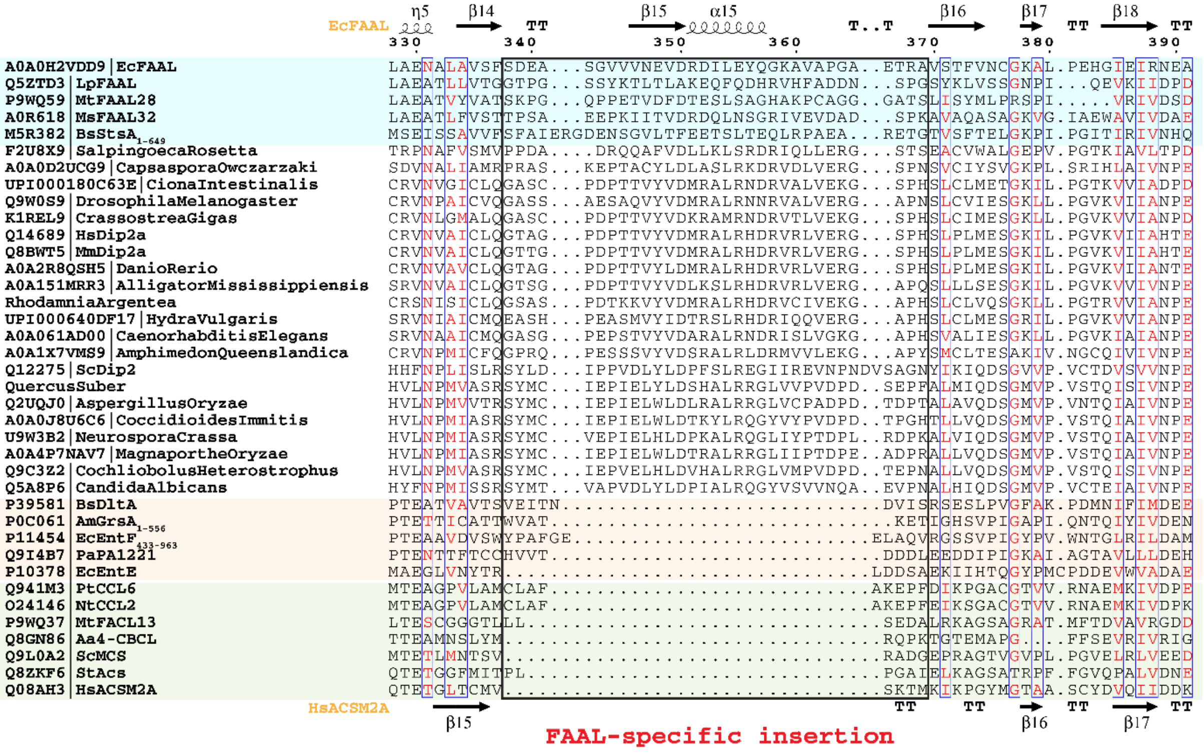

EcFAAL
η5
β14
TT
β15
α15
T..T
β16
β17
TT
β18
TT
330
340
350
360
370
380
390
A0A0H2VDD9|EcFAAL
Q5ZTD3|LpFAAL
P9WQ59|MtFAAL28
A0R618|MsFAAL32
M5R382|BsStsA1-649
F2U8X9|SalpingoecaRosetta
A0A0D2UCG9|CapsasporaOwczarzaki
UPI000180C63E|CionaIntestinalis
Q9W0S9|DrosophilaMelanogaster
K1REL9|CrassostreaGigas
Q14689|HsDip2a
Q8BWT5|MmDip2a
A0A2R8QSH5|DanioRerio
A0A151MRR3|AlligatorMississippiensis
RhodamniaArgentea
UPI000640DF17|HydraVulgaris
A0A061AD00|CaenorhabditisElegans
A0A1X7VMS9|AmphimedonQueenslandica
Q12275|ScDip2
QuercusSuber
Q2UQJ0|AspergillusOryzae
A0A0J8U6C6|CoccidioidesImmitis
U9W3B2|NeurosporaCrassa
A0A4P7NAV7|MagnaportheOryzae
Q9C3Z2|CochliobolusHeterostrophus
Q5A8P6|CandidaAlbicans
P39581|BsDltA
P0C061|AmGrsA1-556
P11454|EcEntF433-963
Q9I4B7|PaPA1221
P10378|EcEntE
Q941M3|PtCCL6
O24146|NtCCL2
P9WQ37|MtFACL13
Q8GN86|Aa4-CBCL
Q9L0A2|ScMCS
Q8ZKF6|StAcs
Q08AH3|HsACSM2A
HsACSM2A
β15
TT
TT
β16
TT
β17
TT
FAAL-specific insertion

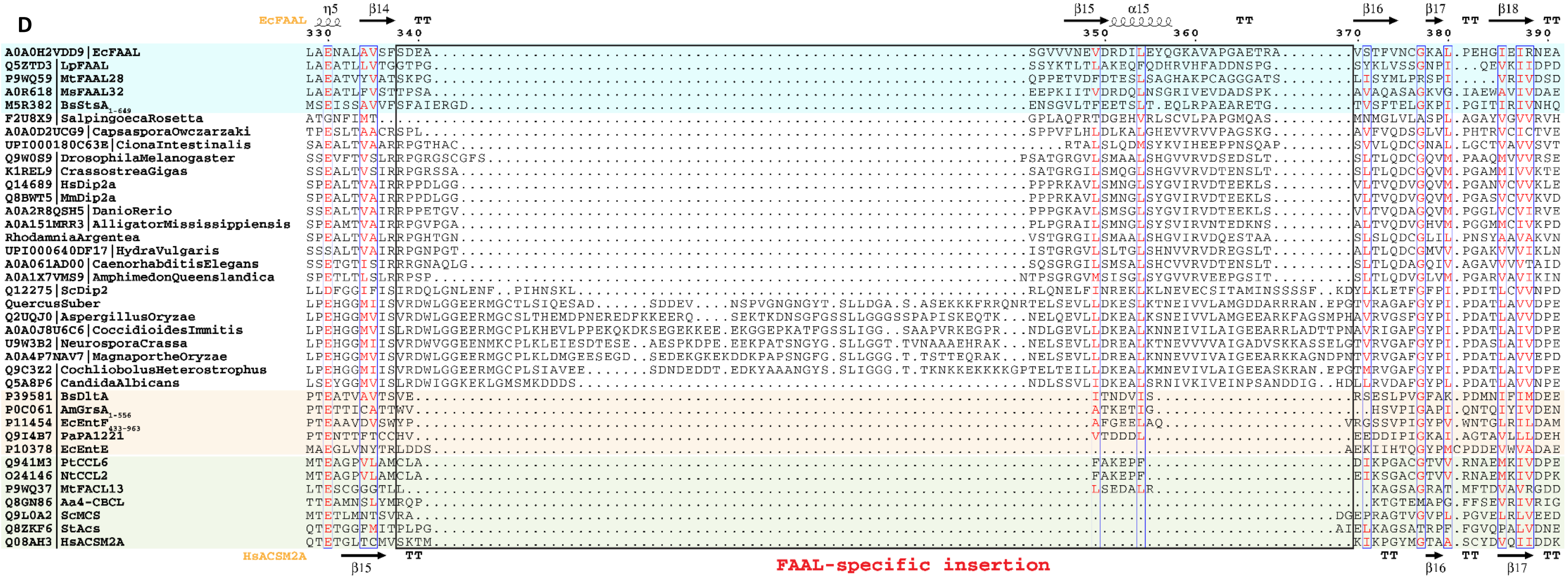
D
EcFAAL
η5
β14
β15
α15
β16
β17
β18
TT
330
340
350
360
370
380
390
A0A0H2VDD9|EcFAAL
Q5ZTD3|LpFAAL
P9WQ59|MtFAAL28
A0R618|MsFAAL32
M5R382|BsStsA1-649
F2U8X9|SalpingoecaRosetta
A0A0D2UCG9|CapsasporaOwczarzaki
UPI000180C63E|CionaIntestinalis
Q9W0S9|DrosophilaMelanogaster
K1REL9|CrassostreaGigas
Q14689|HsDip2a
Q8BWT5|MmDip2a
A0A2R8QSH5|DanioRerio
A0A151MRR3|AlligatorMississippiensis
RhodamniaArgentea
UPI000640DF17|HydraVulgaris
A0A061AD00|CaenorhabditisElegans
A0A1X7VMS9|AmphimedonQueenslandica
Q12275|ScDip2
QuercusSuber
Q2UQJ0|AspergillusOryzae
A0A0J8U6C6|CoccidioidesImmitis
U9W3B2|NeurosporaCrassa
A0A4P7NAV7|MagnaportheOryzae
Q9C3Z2|CochliobolusHeterostrophus
Q5A8P6|CandidaAlbicans
P39581|BsDltA
P0C061|AmGrsA1-556
P11454|EcEntF433-963
Q9I4B7|PaPA1221
P10378|EcEntE
Q941M3|PtCCL6
O24146|NtCCL2
P9WQ37|MtFACL13
Q8GN86|Aa4-CBCL
Q9L0A2|ScMCS
Q8ZKF6|StAcs
Q08AH3|HsACSM2A
HsACSM2A
FAAL-specific insertion

**Supplementary Figure-8:** The FAAL-like sequences were identified and aligned to generate a structure-based alignment as described in the methods sections. The sequences whose structures are available are coloured (FAALs in blue, A-domains in light red and FACLs in green) and the secondary structures of *Ec*FAAL (PDB: 3PBK; Top) along with *Hs*ACSM2A (PDB: 3EQ6) is also included. The FAAL-specific insertion region is marked as a black rectangle and labelled accordingly (red). (**A**) A survey of the various domain organisations of various FAAL-like sequences is depicted as PKS/NRPS-related domain organisations (top left panel) and PKS/NRPS-independent domain organisations (top right panel), while the bottom panels represent rare/miscellaneous domain organisations. The PKS/NRPS-related domain organisations are seen in bacteria, plants and few lower eukaryotes including amoebozoa (e.g: *Acanthamoeba*), Haptista (e.g.: *Emiliania*) and few fungi where bacterial/plant FAAL-like domains (dark green) are associated with a carrier protein domain (violet) and different domains of NRPS/PKS systems (variegated colour). The bacterial/plant FAAL-like domains are occasionally seen in different domain contexts such as alpha aminoadipate reductase domains or α/β-hydrolases (white dotted box) in few bacterial, plant and lower eukaryotic species or sandwiched between HemY (grey) and catalase (olive) domains exclusively found in plants. The PKS/NRPS-independent domain organisations are almost exclusively seen in opisthokonts and rarely in few plants. The FAAL-like sequences of opisthokonts are related to typical bacterial/pPlant FAAL-like sequence but show two different levels of sequence divergence, namely FAAL-like-1 (pale green) and FAAL-like-2 (green), where the lighter shade indicates the extent of divergence. All the identified opisthokonta FAAL-like sequences are always found as a tandem fusion of FAAL-like-1 and FAAL-like-2. This didomain organisation is seen in choanoflagellates (e.g.: *Monosiga*, *Salpingoeca*), while in higher forms it almost exclusively seen as a fusion of didomain architecture with a DMAP1-binding domain (pentagon in light red) forming a highly conserved three-domain architecture. In fungi (and few plant species such as *Quercus* and *Rhodamnia*), the sequence similarity to DMAP1-binding domain is poor, indicated by a serrated pentagon, unlike higher forms. Additional domains such as ARG80, RAMP, PHA03247 family of domains (dotted white box) are occasionally fused to the conserved three-domain architecture. (**B**) A better sequence relationship with FAALs along with subtle differences in various motifs in opisthokont FAAL-like domains is commensurate with the clustering behaviour in the phylogenetic tree generated by different methods. The maximum-likelihood tree implicates a closer relationship of opisthokont FAAL-like domains (pink) to Bacterial FAALs (blue) and Plant FAALs (green) unlike the NJ-tree, nevertheless the partitioning of all FAALs away from A-domains (peach) or FACLs (orange) is noted throughout. **(C)** The FAAL-specific insertion region of FAAL-like-2 shows a comparable length with that of prokaryotic FAALs. **(D)** The FAAL-specific insertion region of FAAL-like-1 in fungi shows an unusually long insertion as compared to the insertion in prokaryotic FAALs, while the higher forms show comparable length.

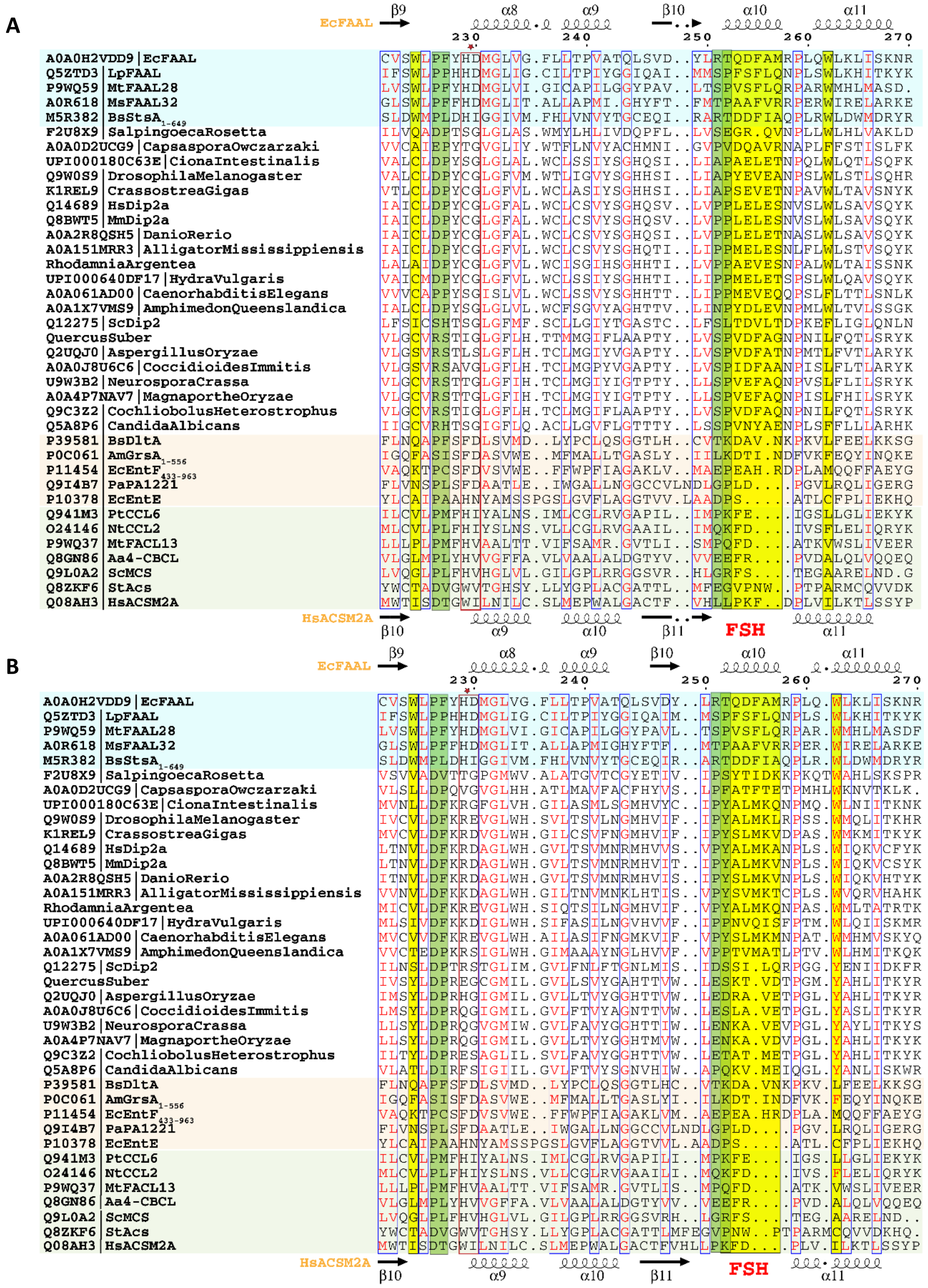

**Supplementary Figure-9:** The FAAL-like sequences were identified and aligned to generate a structure-based alignment as described in the methods sections . The sequences whose structures are available are coloured (FAALs in blue, A-domains in light red and FACLs in green) and the secondary structures of *Ec*FAAL (PDB: 3PBK; Top) along with *Hs*ACSM2A (PDB: 3EQ6) is also included. The FAAL-specific helix (FSH) region is marked as a yellow rectangle and labelled accordingly (red). The bulky residues contributing to the CoA-rejection mechanism are highlighted in yellow. The structural sites that correspond to the prolines of the unique alternative pocket in FAALs are highlighted in green. The A4 motif, which is proposed to harbour the residues important for the catalysis of thioesterification are marked in a red rectangle along with a star **(A)** The FSH region of FAAL-like-2 is a comparable to that of prokaryotic FAALs, while the additional bulky residues such as Trp/Phe (succeeding FSH) is conserved in higher opisthokonts but not fungal FAAL-like-2 sequences whereas the Trp (preceding FSH) is completely absent in both fungi and higher opisthokonts. The prolines of the alternative pocket away from FSH appear in a circular mutation in fungi but in no other opisthokonts, while the prolines of the alternative pocket proximal to FSH appear conserved in all opisthokonts. The A4 motif of the FAAL-like-2 is a conserved Cys in higher opisthokont, however no such conservation in noted in any of the fungal FAAL-like-2 domains. **(B)** The FSH region of FAAL-like-2 is a comparable to that of prokaryotic FAALs, while the additional bulky residues such as Trp/Phe (succeeding FSH) is conserved in higher opisthokonts but not fungal FAAL-like-2 sequences whereas the Trp/Tyr (preceding FSH) is present in fungi but no other opisthokonts. Similar to FAAL-like-2 domains, the prolines of the alternative pocket away from FSH appear in a circular mutation in fungi but no other opisthokonts, while the prolines of the alternative pocket proximal to FSH appear in circular mutation in higher opisthokonts but not fungi. The A4 motif of the FAAL-like-2 is a conserved Arg in higher opisthokont, however, in the fungal FAAL-like-2 domains they may be Gln/Glu/His.

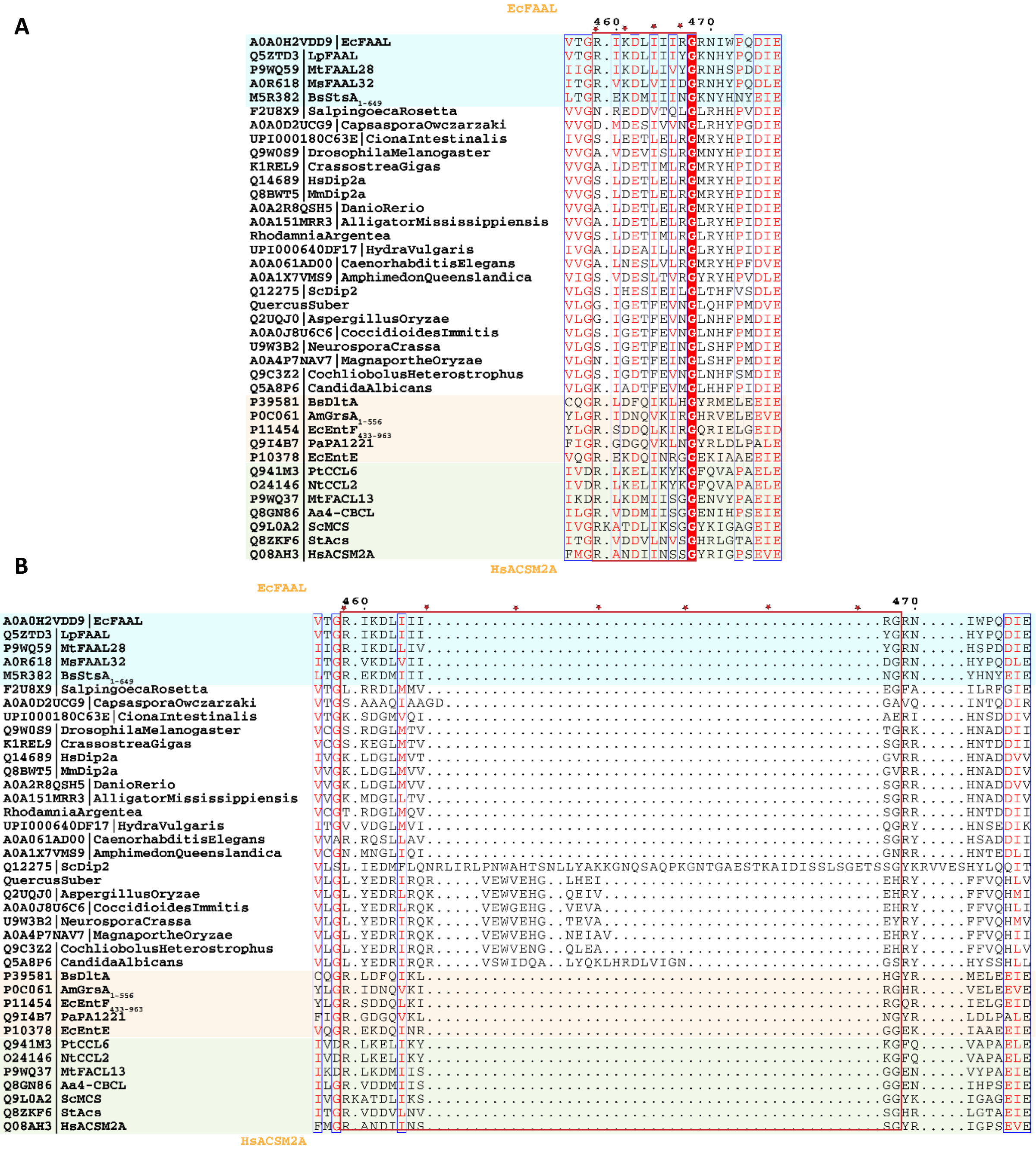
A
EcFAAL
460
470
A0A0H2VDD9|EcFAAL
Q5ZTD3|LpFAAL
P9WQ59|MtFAAL28
A0R618|MsFAAL32
M5R382|BsStsA1-649
F2U8X9|SalpingoecaRosetta
A0A0D2UCG9|CapsasporaOwczarzaki
UPI000180C63E|CionaIntestinalis
Q9W0S9|DrosophilaMelanogaster
K1REL9|CrassostreaGigas
Q14689|HsDip2a
Q8BWT5|MmDip2a
A0A2R8QSH5|DanioRerio
A0A151MRR3|AlligatorMississippiensis
RhodamniaArgentea
UPI000640DF17|HydraVulgaris
A0A061AD00|CaenorhabditisElegans
A0A1X7VMS9|AmphimedonQueenslandica
Q12275|ScDip2
QuercusSuber
Q2UQJ0|AspergillusOryzae
A0A0J8U6C6|CoccidioidesImmitis
U9W3B2|NeurosporaCrassa
A0A4P7NAV7|MagnaportheOryzae
Q9C3Z2|CochliobolusHeterostrophus
Q5A8P6|CandidaAlbicans
P39581|BsDltA
P0C061|AmGrsA1-556
P11454|EcEntF433-963
Q9I4B7|PaPA1221
P10378|EcEntE
Q941M3|PtCCL6
O24146|NtCCL2
P9WQ37|MtFACL13
Q8GN86|Aa4-CBCL
Q9L0A2|ScMCS
Q8ZKF6|StAcs
Q08AH3|HsACSM2A
HsACSM2A
B
EcFAAL
460
470
HsACSM2A

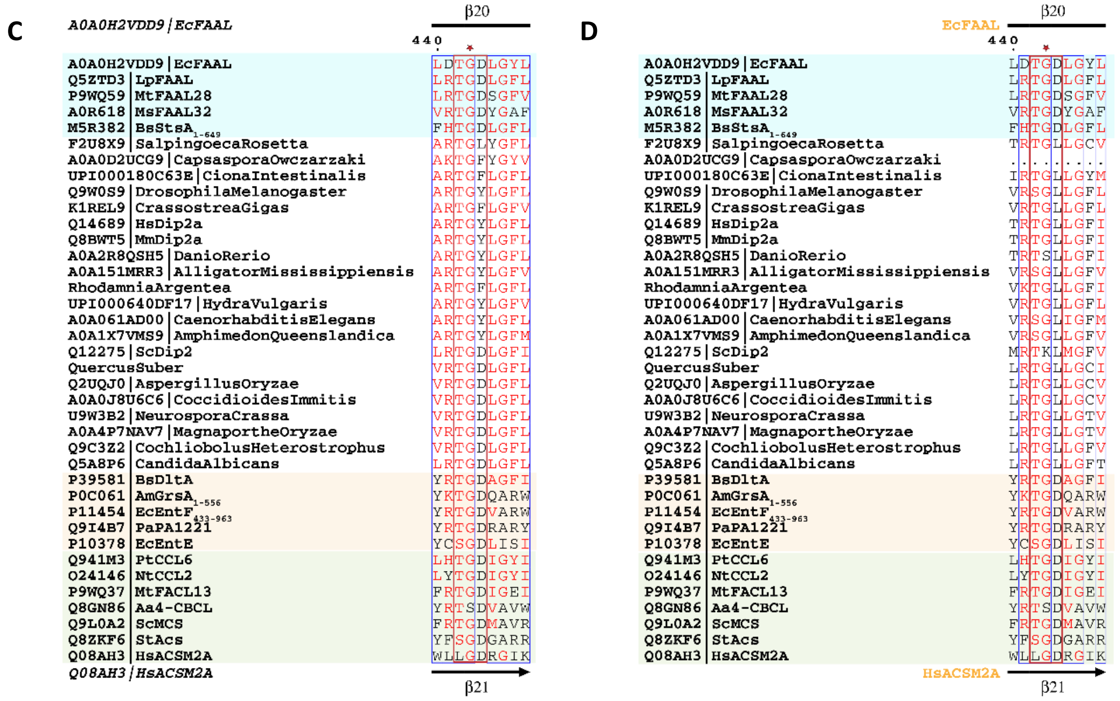

**Supplementary Figure-10:** The FAAL-like sequences were identified and aligned to generate a structure-based alignment as described in the methods sections. The sequences whose structures are available are coloured (FAALs in blue, A-domains in light red and FACLs in green) and the secondary structures of *Ec*FAAL (PDB: 3PBK; Top) along with *Hs*ACSM2A (PDB: 3EQ6) is also included. The A8-motif and A5-motif, which are proposed assist domain alternation and binding ribose of ATP respectively are marked in a red rectangle along with a star(s). The phylogenetic tree was generated from as described in the methods sections. The A8-motif of FAAL-like-2 (**A**) and FAAL-like-1 (**B**) of higher opisthokonts are comparable in length to a typical A8-motif of ANL superfamily unlike the A8-motif of fungal FAAL-like-1 domains. However, in terms of sequence conservation, both FAAL-like-1 or FAAL-like-2 neither resemble FAALs or other members of the superfamily. The A5-motif of fungal FAAL-like-2 (**C**) have a conserved Asp just like all members of the superfamily, but it has been replaced by a conserved Tyr in higher opisthokonts. The A5-motif of all FAAL-like-1 (**D**) have the conserved Asp replaced by a conserved Leu in all opisthokonts.
