## Supplementary table-1 for "A universal pocket in Fatty acyl-AMP ligases ensures redirection of fatty acid pool away from Coenzyme A-based activation"

| Protein | Organism | Uniprot ID | N-terminal only | Undefined | Adenylation state (A-state) | Thioesterification state (T-state) |
|---|---|---|---|---|---|---|
| 2,3-dihydroxybenzoate-AMP ligase | *Acinetobacter baumannii* | A0A0Q1N003 | 3O82, 3O83, 3O84, 3U16, 3U17 | | | |
| ABBFA_003403 | *Acinetobacter baumannii* | A0A0X1KH98 | | 4ZXH, 4ZXI | | |
| 4-Cholrobenzoyl-CoA ligase | *Alcaligenes sp. AL3007* | Q8GN86 | | | 1T5H, 1T5D, 2QVX, 2QVY, 2QVZ, 2QW0, 3CW8, 3DLP | 3CW9 |
| Blue Shifted Luciferase | *Amydetes vivianii* | A0A3F2YLV6 | | 6AAA | | |
| 2-hydroxyisobutyryl-CoA synthetase | *Aquincola tertiaricarbonis* | I3VE75 | | 6HDW, 6HDX, 6HDY, 6HE0, 6HE2 | | |
| 4-coumarate-CoA ligase 1 | *Arabidopsis thaliana* | Q42524 | 3TSY | | | |
| Oxalate-CoA Ligase | *Arabidopsis thaliana* | Q9SMT7 | | | 5IE0 | |
| Long-chain fatty acid-CoA ligase | *Archaeoglobus fulgidus* (strain ATCC 49558) | O30147 | | 3G7S | | |
| D-alanine-D-alanyl carrier protein ligase | *Bacillus cereus* (strain ATCC 14579) | Q81G39 | | | 3DHV, 3FCC, 3FCE | |
| PheA | *Bacillus migulanus* | P0C061 | | | 1AMU | |
| 2-succinyl benzoate-CoA ligase | *Bacillus subtilis* (strain 168) | P23971 | | 5BUQ | 5BUR, 5BUS | 5X8F |
| DhbE | *Bacillus subtilis* (strain 168) | P40871 | | | 1MDF, 1MD9, 1MDB | |
| DltA | *Bacillus subtilis* (strain 168) | P39581 | | | | 3E7W, 3E7X |

| | | | | | | |
|---|---|---|---|---|---|---|
| SrfA-C | *Bacillus subtilis* (strain 168) | Q08787 | | | 2VSQ | |
| Phenylacetate-coenzyme A ligase | *Bacteroides thetaiotaomicron* | Q8AAN6 | | 4RVN | | 4RVO, 4R1M, 4R1L |
| LgrA | *Brevibacillus parabrevis* | Q70LM7 | 5ES6, 5ES7, 5JNF, 6MFY, 6MFZ | 5ES5, 5ES9, 6MFX, 6MFW, 6MG0 | | 5ES8 |
| TycA | Brevibacillus parabrevis | P09095 | 5N81, 5N82 | | | |
| Acyl-CoA Synthetase | *Brucella canis* | A9MB96 | | 5U2A | | |
| Phenylacetate-coenzyme A ligase | *Burkholderia cenocepacia* (strain ATCC BAA-245) | B4E7B5 | | 2Y27, 2Y4N | | |
| Phenylacetate-coenzyme A ligase | *Burkholderia cenocepacia* (strain ATCC BAA-245) | B4EL89 | | | | 2Y4O |
| ObiF1 | *Burkholderia diffusa* | A0A5H1ZR44 | | 6N8E | | |
| Bile acid-coenzyme A ligase | *Clostridium scindens* | P19409 | 4LGC | | | |
| Acetyl-CoA Synthetase | *Cryptococcus neoformans* var. grubii serotype A | J9VFT1 | | 5VPV | 5K85, 5U29, 5K8F, 5IFI, 5VPV | |
| Dshi_0825 | *Dinoroseobacter shibae* | A8LRC0 | | | | 5GXD |
| Acyl-CoA synthetase | *Dyadobacter fermentans* | C6W5A4 | | 4GS5 | | |
| Txo1 | *Eleftheria terrae* | A0A0B5GUD2 | 6OYF, 6OZV, 6P4U | 6P1J | | |
| EhpF | *Enterobacter agglomerans* | Q8GPH0 | 3L2K | | | |
| 3-hydroxyacyl-CoA dehydrogenase | *Erythrobacter sp.* NAP1 | A3WE14 | | | | 6EQO |

| | | | | | | |
|---|---|---|---|---|---|---|
| 2-succinyl benzoate-CoA ligase | *Escherichia coli* (strain K12) | P37353 | | | 5C5H, 6NJ0 | |
| EntE | *Escherichia coli* (strain K12) | P10378 | | | | 6IYK, 6IYL |
| EntE+EntB Fusion | *Escherichia coli* (strain K12) | P0ADI4+P10378 | | | | 3RG2, 4IZ6 |
| EntF | *Escherichia coli* (strain K12) | P11454 | | | | 5T3D, 5JA1, 5JA2 |
| EcFAAL | *Escherichia coli* O6:H1 | A0A0H2VDD9 | | | 3PBK | |
| Nonribosomal peptide synthetase | *Geobacillus sp.* Y4.1MC1 | UPI0001B0E66C | | 5U89 | | |
| ACSM2A | *Homo sapiens* | Q08AH3 | | | 3C5E, 3DAY, 3GPC | 2WD9, 2VZE, 3EQ6, 3B7W |
| AGF18_11095 | *Klebsiella oxytoca* | A0A2U4DY99 | 6VHT, 6VHU, 6VHW, 6VHX, 6VHZ6 | | 6VHV | 6VHY |
| Luciferase | *Lampyris turkestanicus* | Q5UFR2 | 3QYA, 4M46 | | | |
| LpFAAL | *Legionella pneumophila* | Q5ZTD3 | | 3KXW, 3LNV | | |
| Luciferase | *Luciola cruciata* | P13129 | | | 2D1Q, 2D1R, 2D1S, 2D1T | |
| Fatty acid-CoA ligase | *Marinactinospora thermotolerans* | R4R1U5 | | | 6H1B, 6SQ8 | |
| MA_2912 | *Methanosarcina acetivorans* (strain ATCC 35395) | Q8TLW1 | | | | 3ETC |
| McyG | *Microcystis aeruginosa* PCC 7806 | A8YJW1 | | | 4R0M | |
| TioS+TioT fusion | Micromonospora sp. ML1 | Q333U7+Q333U6 | | | | 5WMM |

| | | | | | | |
|---|---|---|---|---|---|---|
| CAR Aden GR01_22995 | *Mycobacterium chelonae* | A0A0E3TT64 | | 6OZ1 | | |
| MsFAAL32 | *Mycobacterium marinum* | B2HMK0 | | | 5EY9 | |
| MbtA | *Mycobacterium smegmatis* | A0R0V0 | | 5KEI | | |
| MsFAAL32 | *Mycobacterium smegmatis* | A0R618 | 5D6N | | 5D6J, 5EY8, 5ICR | |
| FAAL28 | *Mycobacterium tuberculosis* H37Rv | P9WQ59 | 3E53, 3T5A | | | |
| FACL13 | *Mycobacterium tuberculosis* H37Rv | P9WQ37 | 3T5B, 3T5C, 5ZRN | 3R44 | | |
| FadD10 | *Mycobacterium tuberculosis* H37Rv | P9WQ55 | | 4IR7, 4ISB | | |
| FadD32 | *Mycobacterium tuberculosis* H37Rv | O53580 | | | 5HM3 | |
| Nonribosomal peptide synthase SidN | *Neotyphodium lolii* | K7NCP5 | 3ITE | | | |
| 4-coumarate-CoA ligase 2 | *Nicotiana tabacum* | O24146 | | 5U95 | 5BSM, 5BSW | 5BSR, 5BST, 5BSU, 5BSV |
| Carboxylic Acid Reductase | *Nocardia iowensis* | Q6RKB1 | | 5MSC, 5MSD | | |
| Benzoate-CoA ligase | *Paraburkholderia xenovorans* (strain LB400) | Q13WK3 | | | 2V7B | |
| Luciferase | *Photinus pyralis* | P08659 | 3IEP, 3IER, 3IES, 3RIX | 1LCI, 1BA3, 5DV9 | 4G36, 6SQ8, 5KYT, 5KYV, 6Q2M | 4G37 |
| ApnA A1 | *Planktothrix rubescens* NIVA-CYA 18 | G0WVH3 | 4D4H | | | 4D4G, 4D4I, 4D56, 4D57 |
| 4-coumarate:CoA ligase | *Populus tomentosa* | Q941M3 | | 3A9U | | 3A9V |

| | | | | | | |
|---|---|---|---|---|---|---|
| PA0996 | *Pseudomonas aeruginosa PAO1* | Q9I4X3 | 5OE3, 5OE4, 5OE5, 5OE6 | | | |
| PA1221 | *Pseudomonas aeruginosa* PAO1 | Q9I4B7 | | | | 4DG8, 4DG9 |
| PltF+PltL coupled | *Pseudomonas fluorescens* | Q4KCY5+Q4KCZ1 | | | | 6O6E |
| Benzoate-CoA Ligase | *Rhodopseudomonas palustris* | Q93TK0 | | | | 4EAT, 4RLF, 4RLQ, 4RM2, 4RM3, 4RMN, 4ZJZ |
| Long-chain fatty acid-CoA ligase | *Rhodopseudomonas palustris* | Q6NAE3 | 3IVR | | | |
| Malonyl CoA synthetase | *Rhodopseudomonas palustris* | Q6ND88 | | 4FUQ, 4FUT, 4GXQ, 4GXR | | |
| SL1157_2728 | *Ruegeria lacuscaerulensis* | D0CPY8 | | 6IHK, 6IJB | | |
| Acetyl-CoA synthetase | *Saccharomyces cerevisiae* | Q01574 | | | 1RY2 | |
| Acetyl-CoA synthetase | *Salmonella typhimurium* | Q8ZKF6 | | | | 1PG3, 1PG4, 2P2F, 2P20, 2P2B, 2P2J, 2P2M, 2P2Q, 5JRH |
| Carboxylic Acid Reductase | *Segniliparus rugosus* ATCC BAA-974 | E5XP76 | | 5MST, 5MSW | | 5MSS |
| 2-succinyl benzoate-CoA ligase | *Staphylococcus aureus* (strain N315) | P63526 | | 3IPL | | |
| Anthranilate-CoA ligase | *Stigmatella aurantiaca* | F3Y661 | | | 4WV3 | |
| D-alanine-D-alanyl carrier protein ligase | *Streptococcus pyogenes* serotype M6 | Q5XBN5 | | 3L8C, 3LGX | | |
| Malonyl-CoA Ligase | *Streptomyces coelicolor* (strain ATCC BAA-471) | Q9L0A2 | | | | 3NYQ, 3NYR |

| | | | | | | |
|---|---|---|---|---|---|---|
| CahJ | Streptomyces gandocaensis | A0A140DJY3 | | | | 5WM2, 5WM3, 5WM4, 5WM5, 5WM6, 5WM7 |
| VinN | *Streptomyces halstedii* | Q76KY2 | 3WVN, 3WV5, 3WV4 | | | |
| Acetoacetyl-CoA Synthetase 1 | *Streptomyces lividans* TK24 | D6EQU8 | | | 4WD1 | |
| PtmA2 | *Streptomyces platensis* | | | 5E7Q, 5UPQ | | |
| CmiS6 | *Streptomyces* sp MJ635-86F5 | X5IJ97 | 5JJP | | | |
| Thr1 | *Streptomyces sp.* | H6SG27 | | | 5N9X, 5N9W | |
| IdnL1 | *Streptomyces* sp. ML694-90F3 | A0A077KT11 | | | | 5JJQ |
| IdnL7 | *Streptomyces sp.* ML694-90F3 | A0A077KUW8 | | 6AKD | | |
| CytC1 | *Streptomyces* sp. NBRC 110028 | UPI0006E2E4D2 | | 3VNR, 3VNS, 3VNQ | | |
| Fatty acyl-CoA synthetase | *Thermus thermophilus* (strain HB8) | Q5SKN9 | | 1ULT | | 1V25, 1V26 |
| EntF | *Vibrio cholerae* serotype O1 | Q9KRQ7 | | | | 4OXI |
| Luciferase-like enzyme AMP-CoA-ligase | *Zophobas atratus* | A0A0R4I967 | 4W8O | | | |

Supplementary Table-I: List of PDBs of the ANL superfamily family members curated from the RCSB-PDB database.
