## Supplementary table-2 for "A universal pocket in Fatty acyl-AMP ligases ensures redirection of fatty acid pool away from Coenzyme A-based activation"

| | Number of atoms in FAALs at ≤ 2.5 Å from multiple CoA conformations | | | | | |
|---|---|---|---|---|---|---|
| | 1PG4_CoA | 3EQ6_CoA | 3CW9_CoA | 5BSR_CoA | 3NYQ_CoA | Average |
| 3PBK | 9 | 9 | 5 | 13 | 2 | 7.6 |
| 5ICR | 8 | 9 | 10 | 10 | 2 | 7.8 |
| 3E53 | 5 | 5 | 9 | 6 | 2 | 5.4 |
| 3KXW | 7 | 4 | 10 | 8 | 2 | 6.2 |
| | | | | | | 7 |

| | Number of atoms in FACLs at ≤ 2.5 Å from multiple CoA conformations | | | | | |
|---|---|---|---|---|---|---|
| | 1PG4_CoA | 3EQ6_CoA | 3CW9_CoA | 5BSR_CoA | 3NYQ_CoA | Average |
| 1PG4 | 0 | 0 | 5 | 3 | 0 | 1.6 |
| 3EQ6 | 4 | 0 | 4 | 3 | 0 | 2.2 |
| 3CW9 | 6 | 0 | 0 | 1 | 0 | 1.4 |
| 5BSR | 5 | 1 | 1 | 0 | 0 | 1.4 |
| 3R44 | 3 | 0 | 0 | 0 | 1 | 0.8 |
| 3A9V | 7 | 0 | 0 | 0 | 0 | 1.4 |
| 3NYQ | 8 | 1 | 2 | 3 | 0 | 2.8 |
| 4DG9 | 0 | 0 | 1 | 0 | 0 | 0.2 |
| | | | | | | 1 |

| | Number of atoms in FAALs at ≤ 2.0 Å from multiple CoA conformations | | | | | |
|---|---|---|---|---|---|---|
| | 1PG4_CoA | 3EQ6_CoA | 3CW9_CoA | 5BSR_CoA | 3NYQ_CoA | Average |
| 3PBK | 1 | 7 | 2 | 7 | 2 | 3.8 |
| 5ICR | 4 | 3 | 5 | 6 | 1 | 3.8 |
| 3E53 | 2 | 4 | 6 | 4 | 2 | 3.6 |
| 3KXW | 3 | 1 | 6 | 6 | 1 | 3.4 |
| | | | | | | 4 |

| | Number of atoms in FACLs at ≤ 2.0 Å from multiple CoA conformations | | | | | |
|---|---|---|---|---|---|---|
| | 1PG4_CoA | 3EQ6_CoA | 3CW9_CoA | 5BSR_CoA | 3NYQ_CoA | Average |
| 1PG4 | 0 | 0 | 3 | 2 | 0 | 1 |
| 3EQ6 | 3 | 0 | 3 | 2 | 0 | 1 |
| 3CW9 | 5 | 0 | 0 | 0 | 0 | 1 |
| 5BSR | 4 | 0 | 1 | 0 | 0 | 1 |
| 3R44 | 1 | 0 | 0 | 0 | 0 | 0 |
| 3A9V | 4 | 0 | 0 | 0 | 0 | 1 |
| 3NYQ | 4 | 0 | 0 | 1 | 0 | 1 |
| 4DG9 | 3 | 0 | 1 | 0 | 0 | 0 |
| | | | | | | 1 |

Supplementary Table-II: A tabulation of the number of atoms from the N-terminal domain of FAALs and FACLs (excluding hydrogens) at a defined distance from the atoms in multiple conformations of CoA seen in the CoA-bound structures of FACLs. The number of atoms of the protein at a clashing distance is an indicator of the space available in the canonical pocket. A higher number of atoms in FAALs indicate the limited space available in the pocket, while the lower number indicates that it is more accommodative in the case of FACLs. Crystal structures of known FACLs have at least one atom clashing with any of the known CoA conformations, while FAALs clash with an average of 7 atoms at 2.5 Å (4 atoms at 2.0Å). Therefore, FAALs have atleast 4-fold greater number of atoms in the vicinity of CoA than FACLs, making FAALs less accommodative of any known CoA conformation from ANL superfamily as compared to FACLs.
