## Supplementary table-3 for "A universal pocket in Fatty acyl-AMP ligases ensures redirection of fatty acid pool away from Coenzyme A-based activation"

| **Lineage** | **Organisms** | **Type of FAAL-like domain (domain organization)** |
|---|---|---|
| Amoebozoa | L8HIJ4\|*Acanthamoeba castellanii* | Bacterial/Plant FAAL |
| Ancyromonadida | - | - |
| Apusozoa | - | - |
| Breviatea | - | - |
| CRuMs | - | - |
| Cryptophyceae | - | - |
| Discoba | - | - |
| Glaucocystophyceae | - | - |
| Haptista | R1DLD1\|*Emiliania huxleyi* | Bacterial/Plant FAAL |
| Hemimastigophora | - | - |
| Malawimonadida | - | - |
| Metamonada | - | - |
| Opisthokonta | | |
| • Aphelida | - | - |
| • Choanoflagellata | A9UP98\|*Monosiga brevicollis*; F2U8X9\|*Salpingoeca rosetta* | Opisthokonta FAAL-like domains (two-domain) |
| • Filasterea | A0A0D2UCG9\|*Capsaspora owczarzaki* | Opisthokonta FAAL-like domains (three-domain) |
| • Fungi | Many organisms (except Basidiomycetes) | Opisthokonta FAAL-like domains (three-domain) |
| • Ichthyosporea | - | - |
| • Metazoa | | Opisthokonta FAAL-like domains (three-domain) |
| • Rotosphaerida | - | - |
| Rhodelphea | - | - |
| Rhodophyta | - | - |
| SAR group | | |
| • Stramenopiles | A0A067C5L3\|*Saprolegnia parasitica*; A0A1V9Z9Z4\|*Achlya hypogyna* | Bacterial/Plant FAAL |
| • Alveolata | V4ZF64\|Toxoplasma gondii; U6GWK7\|*Eimeria acervuline* | Bacterial/Plant FAAL |
| • Rhizaria | - | - |
| Viridiplantae | A0A445HLB4\|*Glycine soja*; A0A3Q0EXH4\|*Vigna radiata*<br>Q01KB0 \|*Oryza sativa*;<br>UPI000CE285CA\|*Quercus suber*; UPI0011E53342\|*Rhodamnia argentea* | Bacterial/Plant<br>Bacterial/Plant fused to HemY/Catalase<br>Di-domain form (tandem FAAL-like) |

**Supplementary Table-III:** A tabulation of the various FAAL-like domains and their domain organisations identified in the different lineages of eukaryotes as per the taxonomic distribution provided at NCBI Taxonomy (url: http://www.ncbi.nlm.nih.gov/taxonomy) [1].

1 Schoch, C. L. *et al.* NCBI Taxonomy: a comprehensive update on curation, resources and tools. *Database (Oxford)* **2020**, doi:10.1093/database/baaa062 (2020).
