## Supplementary table-4 for "A universal pocket in Fatty acyl-AMP ligases ensures redirection of fatty acid pool away from Coenzyme A-based activation"

Supplementary table-IV: Primers used for cloning and generating mutants of proteins in this study.

EcFAAL (A0A0H2VDD9): Cloned as a hexahistidine-tagged protein fused to SUMO tag at the N-terminus
5' GCAGCCATATGTCTAATAAAATCTTTACGCATTCCC 3'
5' CCAGACTCGAGTTATGCCAGGGATTCCTGCACATTA 3'

Canonical pocket mutants of EcFAAL

1. A253G EcFAAL
   5' CTCAGGATTTTGGCATGCGTCCTCTGCAATGGC 3'
   5' AGAGGACGCATGCCAAAATCCTGAGTGCGCAAA 3'
2. F275A EcFAAL
   5' GTTGCGCCGCCGGCGGGCTATGAATTGTGCCAGCG 3'
   5' CAATTCATAGCCCGCCGGCGGCGCAACGGAAACGG 3'
3. M227A
   5' TTCTACCATGATGCGGGACTGGTCGGCTTTCTCC 3'
   5' CCGACCAGTCCCGCATCATGGTAGAAAGGCAGCC 3'
4. ΔFSH (DFAM)
   5' GCTTTCAGTAGATTATTTGCGCACTCAGCGTCCTCTGCAATGGCTTAAATTGATCAGTAAAAATC 3'
   5' CTGATCAATTTAAGCCATTGCAGAGGACGCTGAGTGCGCAAATAATCTACTGAAAGCTGCGTGGC 3'
5. ΔFSI
   5' GGTTGTGGTTAACGAAGTGGATCCGGGTGCAGAGACACGCGCCGTATC 3'
   5' GCGTGTCTCTGCACCCGGATCCACTTCGTTAACCACAACCCCGGAGGC 3'

Alternative pocket mutants of EcFAAL

1. T83F EcFAAL
   5' CTGATTGCCGAATTTAGTAGCGAGTTCGTAGAGGC 3'
   5' GAACTCGCTACTAAATTCGGCAATCAGTGCCACGC 3'
2. T83R EcFAAL
   5' CTGATTGCCGAACGCAGTAGCGAGTTCGTAGAGGC 3'
   5' GAACTCGCTACTGCGTTCGGCAATCAGTGCCACGC 3'
3. P107F
   5' CCGTTGGCGATTTTTATGGGCGTTGGTCAGCGGGA 3'
   5' ACCAACGCCCATAAAAATCGCCAACGGGACGGCGA 3'
4. P107R
   5' CGTTGGCGATTCGCATGGGCGTTGGTCAGCGGGA 3'
   5' ACCAACGCCCATGCGAATCGCCAACGGGACGGCG 3'
5. P107L
   5' CGTTGGCGATTCTGATGGGCGTTGGTCAGCGGGA 3'
   5' ACCAACGCCCATCAGAATCGCCAACGGGACGGCG 3'
6. T252F
   5' GTCTCCTGGCTGTTTTTCTACCATGATATGGGAC 3'
   5' TCATGGTAGAAAAACAGCCAGGAGACGCAGCGGT 3'
7. T252R
   5' TCTCCTGGCTGCGCTTCTACCATGATATGGGACT 3'
   5' ATCATGGTAGAAGCGCAGCCAGGAGACGCAGCGG 3'

EcACP (A0A2X1NC35): Cloned as a hexahistidine-tagged protein at the C-terminus
5' GCAGCCATATGGTAAATCGTGAAATAGTAATG 3'
5' GTGGTGCTCGAGTTTATTCTCCAGCCATGG 3'

S39A EcACP (apo mutant)
5' CAACGATCTTGGCCTGGAGGCGATCAAAGTGATGGATCTCTTAATGATGCTGGAAG 3'
5' CATCATTAAGAGATCCATCACTTTGATCGCCTCCAGGCCAAGATCGTTGACGAGATC 3'

MsFAAL32 (A0R618): Cloned as a hexahistidine-tagged protein fused to SUMO tag at the N-terminus

| |
|---|
| 5' GCGTCTGAGCTCATGCCGTTCCACAATCCGTTCATC 3'<br>5' GTGGCGGCCGCTCTATTAGTCGGTGG 3' |
| Canonical pocket mutants of MsFAAL32<br>1. A253G MsFAAL32<br>5' CCGGCCGCGTTCGCGCGTCGCCCCGAGCGCTGGATC 3'<br>5' GCTCGGGGCGACGCGCGAACGCGGCCGGGGTCATGA 3'<br>2. F284A MsFAAL32<br>5' GGTGGCCCCGAACGCTGCGTTCGACCACGCCGCG 3'<br>5' CGTGGTCGAACGCAGCGTTCGGGGCCACCGAGAT 3'<br>3. M233A MsFAAL32<br>5' TTCTTCCACGACGCGGGTCTGATCACCGCGCTGC 3'<br>5' GTGATCAGACCCGCGTCGTGGAAGAACGGCAGCC 3'<br>4. ΔFSH MsFAAL32<br>5' CCTTCATGACCCCGGCCCGCCCCGAGCGCTGGATCCGCG 3'<br>5' CCAGCGCTCGGGGCGGGCCGGGGTCATGAAGGTGAAGTAG 3'<br>5. ΔFSI MsFAAL32<br>5' CCGAAGATCATCACGGTCGACGACGCCGATTCGCCCAAGGCCGTC 3'<br>5' CTTGGGCGAATCGGCGTCGTCGACCGTGATGATCTTCGGCTCTTCC 3' |
| Alternative pocket mutants of MsFAAL32<br>1. P85F MsFAAL32<br>5' GCCATCCTGTGTTTTCAGAACCTCGATTACCTGGT 3'<br>5' ATCGAGGTTCTGAAAACACAGGATGGCCACACGGT 3'<br>2. P85R MsFAAL32<br>5' CCATCCTGTGTCGCCAGAACCTCGATTACCTGG 3'<br>5' TCGAGGTTCTGGCGACACAGGATGGCCACACGG 3'<br>3. P110F MsFAAL32<br>5' CCGCTGTTCGATTTTTCCGAGCCCGGCCACGTC 3'<br>5' CCGGGCTCGGATTTATCGAACAGCGGCACCGC 3'<br>4. P110R MsFAAL32<br>5' CCGCTGTTCGATCGCTCCGAGCCCGGCCACGTC 3'<br>5' CCGGGCTCGGAGCGATCGAACAGCGGCACCGC 3'<br>5. P113F MsFAAL32<br>5' GATCCGTCCGAGTTTGGCCACGTCGGCCGCCTGCA 3'<br>5' GCCGACGTGGCCAAACTCGGACGGATCGAACAGCG 3'<br>6. P113R MsFAAL32<br>5' GCTGTTCGATCCGTCCGAGCGCGGCCACGTCGGCCGCCTGC 3'<br>5' GCCGACGTGGCCGCGCTCGGACGGATCGAACAGCGGCACCG 3'<br>7. P253F MsFAAL32<br>5' CACCTTCATGACCTTTGCCGCGTTCGTCCGTCGC 3'<br>5' CGGACGAACGCGGCAAAGGTCATGAAGGTGAAGTA 3'<br>8. P253R MsFAAL32<br>5' CCTTCATGACCCGCGCCGCGTTCGTCCGTCGCCC 3'<br>5' GACGAACGCGGCGCGGGTCATGAAGGTGAAGTAG 3' |
| MsPKS13$_{1-1042}$ (A0R617): Residues 1-1042 (excludes the terminal ACP and thioesterase domains) was cloned as a hexahistidine-tagged protein as a GFP fusion at the C-terminus<br>5' GAGATCATATGACCGTCAACGAGATGCGG 3'<br>5' GCACCAGCTAGCGAACAGCGTGCGGAAGTCCAG 3' |
| S38A MsPKS13$_{1-1042}$ (apo mutant)<br>5' CGAACTCGGCCTGTCCGCGCGCGATGCGGTCGCGATGGCCAGC 3'<br>5' CATCGCGACCGCATCGCGCGCGGACAGGCCGAGTTCGACCATCGG 3' |

MxFAAL (Q1CXX0): Cloned as a hexahistidine-tagged protein at the N-terminus
5' GCGGAATTCCATATGAAGGGCGCTGGTTCGCGACTG 3'
5' GCGAAGCTTAGATCTCGCCGTGCTCGGCCCCACG 3'

Canonical pocket mutants of MxFAAL

1. L256G MxFAAL
   5' GATTCCCCCGAGGCCTTCGGCGCACGGCCCGCGTTGTGGCTCCG 3'
   5' CAACGCGGGCCGTGCGCCGAAGGCCTCGGGGGAATCAGCACCAG 3'
2. F279A MxFAAL
   5' CCCGCGCCGAACGCGGCCTATGGCCTGTGTTTGAAG 3'
   5' ACAGGCCATAGGCCGCGTTCGGCGCGGGAGAGATG 3'
3. M231A MxFAAL
   5' CTGTACCACGACGCGGGGCTCATCGGCTGTCTGC 3'
   5' CCGATGAGCCCCGCGTCGTGGTACAGCGGCACCG 3'
4. ΔFSH MxFAAL (AFLA)
   5' GTGCTGATTCCCCCCGAGCGGCCCGCGTTGTGGCTCCGCGC 3'
   5' GCCACAACGCGGGCCGCTCGGGGGGAATCAGCACCAGGCTTC 3'
5. ΔFSI MxFAAL
   5' CGCGCTGGGCGTGGACGGTTCGCGTGAGTTGGTGAGCGTGGGC 3'
   5' CTCACCAACTCACGCGAACCGTCCACGCCCAGCGCGCGCGGCC 3'

Alternative pocket mutants of MxFAAL

1. T83F MxFAAL
   5' GTTGCTGCTGCCCTTTTCGCCGGCCTTCATGGATGC 3'
   5' GAAGGCCGGCGAAAAGGGCAGCAGCAACGCCACCC 3'
2. T83R MxFAAL
   5' GTTGCTGCTGCCCCCGTCGCCGGCCTTCATGGATG 3'
   5' GAAGGCCGGCGACGGGGGCAGCAGCAACGCCACCCG 3'
3. P106F MxFAAL
   5' GTGCCGCTGTACTTTCCGGTGCGGCTGGGGCGACTG 3'
   5' CCAGCCGCACCGGAAAGTACAGCGGCACCGGCACCG 3'
4. P106R MxFAAL
   5' GTGCCGCTGTACCGCCCGGTGCGGCTGGGGCGACTG 3'
   5' CCAGCCGCACCGGGCGGTACAGCGGCACCGGCACC 3'
5. P107F MxFAAL
   5' CCGCTGTACCCGTTTGTGCGGCTGGGGCGACTGGA 3'
   5' CCCCAGCCGCACAAACGGGTACAGCGGCACCGGCA 3'
6. P107R MxFAAL
   5' CGCTGTACCCGCGCGTGCGGCTGGGGCGACTGGA 3'
   5' CCCCAGCCGCACGCGCGGGTACAGCGGCACCGGC 3'
7. P252F MxFAAL
   5' GTGCTGATTCCCTTTGAGGCCTTCCTCGCACGGCC 3'
   5' GCGAGGAAGGCCTCAAAGGGAATCAGCACCAGGCTTCCCGGGTAG 3'
8. P252R MxFAAL
   5' GCCTGGTGCTGATTCCCCGCGAGGCCTTCCTCGCACGGCCCGC 3'
   5' GTGCGAGGAAGGCCTCGCGGGGAATCAGCACCAGGCTTCCCGGG 3'

MxACP (A0R617)
Cloned as a hexahistidine-tagged protein at the C-terminus
5' GCGGAATTCCATATGACGGAGATTCATCGCATCGTCAC 3'
5' GCGGGATCCGTCGCGAACCAGCGCCCTTCA 3'

S38A MxACP (apo mutant)
5' GGACCTCCAGTTGGACGCGCTGGGCCTCACGGTGCTGGCGGTGGG 3'
5' AGCACCGTGAGGCCCAGCGCGTCCAACTGGAGGTCCCGCACCAG 3'

| |
|---|
| RsFAAL (Q8XRP4): Cloned as a hexahistidine-tagged protein fused to SUMO-tag at the N-terminus<br>5' CCGAATGAGCTCATGACCGCCAGTCGTGCCG 3'<br>5' AGCGGTCTCGAGTTAGGCAACCGGTTCCGGTTCC 3' |
| Canonical pocket mutants of RsFAAL<br>1. I241G RsFAAL:<br>5' CTGCTGAGTCCGCTGCTGTTTGGCCAGAAACCGGTTCGTTGGCTGCGTCTGATTAG 3'<br>5' CAGCCAACGAACCGGTTTCTGGCCAAACAGCAGCGGACTCAGCAGGGTAATCGG 3'<br>2. F264A RsFAAL<br>5' CACCAGCGGTGGCCCGAATGCGGGTTATCAGGCCTGTGTGGAACGCG 3'<br>5' CACACAGGCCTGATAACCCGCATTCGGGCCACCGCTGGTGGTAATGCG 3'<br>3. M261A RsFAAL<br>5' GTGGCTGCCTCCGTATCATGATGCGGGCCTGATTGGCGGCGTTCTGAGTCC 3'<br>5' GAACGCCGCCAATCAGGCCCGCATCATGATACGGAGGCAGCCACACCATGCC 3'<br>4. ΔFSH RsFAAL (LFIQ)<br>5' GACGCAGCCAACGAACCGGTTTCAGCGGACTCAGCAGGGTAATCGGA 3'<br>5' CCCTGCTGAGTCCGCTGAAACCGGTTCGTTGGCTGCGTCTGATTAG 3'<br>5. ΔFSI RsFAAL<br>5' GGCGCCGGTTATAAAGTGCTGTTTCGCGATGCAGTTAGCTGTGGCAGTGCAC 3'<br>5' CCACAGCTAACTGCATCGCGAAAGCACTTTATAACCGGCGCCGCGGGCAC 3' |
| Alternative pocket mutants of RsFAAL<br>1. A71F RsFAAL<br>5' GAACGTGTGCTGATTCTGTGTCGCGTGGGTGAAGAATATACCAGCGTTTTCTTTGC 3'<br>5' CGCTGGTATATTCTTCACCCACGCGACACAGAATCAGCACACGTTCGCGCTGCA 3'<br>2. A71R RsFAAL<br>5' 3'<br>5' 3'<br>3. P95F RsFAAL<br>5' GTTGCCGTTCCGTGTACCTTTCTGCGCGGCATGCGCCATCGCAC 3'<br>5' GGCGCATGCCGCGCAGAAAGGTACACGGAACGGCAACTGCACCGG 3'<br>4. P95R RsFAAL<br>5' CAGTTGCCGTTCCGTGTACCCGCCTGCGCGGCATGCGCCATCGCAC 3'<br>5' GCGCATGCCGCGCAGGCGGGTACACGGAACGGCAACTGCACCGGCC 3'<br>5. P237F RsFAAL<br>5' GTTATCCGATTACCCTGCTGAGTTTTCTGCTGTTTATTCAGAAACCGGTTCGTTGGC 3'<br>5' CCGGTTTCTGAATAAACAGCAGAAAACTCAGCAGGGTAATCGGATAACCGGTATAA 3'<br>6. P237R RsFAAL<br>5' GTTATCCGATTACCCTGCTGAGTCGCCTGCTGTTTATTCAGAAACCGGTTCGTTGGC 3'<br>5' CCGGTTTCTGAATAAACAGCAGGCGACTCAGCAGGGTAATCGGATAACCGGTATAA 3' |
| RsACP (Q8XRP0)<br>Cloned as a hexahistidine-tagged protein at the C-terminus<br>5' GAAGGAGCATATGAACCAGCCGCCGAG 3'<br>5' GAAGTACCTCGAGTGCTTCACTACCGG 3' |
| S48A RsACP (apo mutant)<br>5' GCCGATCATGGCGTTGATGCGATGATTGCCATTGTTATGAGTGGTGACCTGAGC3'<br>5' CACCACTCATAACAATGGCAATCATCGCATCAACGCCATGATCGGCAAAGGTGG 3' |

MtFACL13 (P9WQ37): Cloned as a hexahistidine-tagged protein at the N-terminus
5' CATCATCACCACCATCACAAGAACATTGGCTGGATGCTCAGAC 3'
5' GTGGCGGCCGCTCTATTACTTCGGCACCGTCGCCGAATAC 3'

Canonical pocket mutants of MtFACL13

1. A253F MtFACL13
   5' GGTGGCGCCGTGCCGTTCATCCTCAACTTCATGCGCCAGGTG 3'
   5' CATGAAGTTGAGGATGAACGGCACGGCGCCACCGATACAGAC 3'
2. A210M MtFACL13
   5' ATGTTCCACGTGATGGCGTTGACGACGGTCATCT 3'
   5' GTCGTCAACGCCATCACGTGGAACATCGGCAGCG 3'
3. $FSH_{ins}$ MtFACL13 (SFQL)
   5' ATGCCGCAGTTCAGCTTTCTGCAGGATGCGACGAAGGTGTGGTC 3'
   5' CTTCGTCGCATCCTGCAGAAAGCTGAACTGCGGCATCGAGATCAG 3'

AfFACL (P9WQ37): Cloned as a hexahistidine-tagged protein at the N-terminus
5' GGCAGCCATATGGAGTTGAAGTACAAAATCGGA 3'
5' CGTCCGTCGACACCCTTTTCGGCCTCCTTCT 3'

Canonical pocket mutants of AfFACL

1. A276F AfFACL
   5' GCCGTTCCTCCATTTCTCAACGTGCTCGTCAACAC 3'
   5' GAGCACGTTGAGAAATGGAGGAACGGCCCATGAAA 3'
2. A232M AfFACL
   5' ATGTTCCATTCAATGGAGTTTGGACTGGTCAATT 3'
   5' AGTCCAAACTCCATTGAATGGAACATTGGCATGC 3'
3. $FSH_{ins}$ AfFACL (SFQL)
   5' GGAATGTTTAACAGCTTTCTGCAGCAGGAGATGCTGGCTGAGAA 3'
   5' CAGCATCTCCTGCTGCAGAAAGCTGTTAAACATTCCCATCACAA 3'

EcFACL (P69451): Cloned as a hexahistidine-tagged protein at the C-terminus
5' CATATGGCTAGCATGAAGAAGGTTTGGCTTAACCGTTA 3'
5' GTGGTGCTCGAGGGCTTTATTGTCCACTTTGC 3'

Canonical pocket mutants of EcFACL

1. T308F EcFACL
   5' ACGGGCGTTAACTTTTTGTTCAATGCGTTGCTGAA 3'
   5' CGCATTGAACAAAAAGTTAACGCCCGTGATAGCGG 3'
2. A264M EcFACL
   5' GCTGCCGCTGTATCACATTTTTATGCTGACCATTAACTGCCTGCTGTTTATCGAAC 3'
   5' CAGCAGGCAGTTAATGGTCAGCATAAAAATGTGATACAGCGGCAGCGCCGTCAC 3'
3. $FSH_{ins}$ EcFACL (SFQL)
   5' GCTTATCACTAACCCGCGCAGCTTTCTGCAGGATATTCCAGGGTTGGTAAAAGAGTTAGC 3'
   5' CTTTTACCAACCCTGGAATATCCTGCAGAAAGCTGCGCGGGTTAGTGATAAGCAGGTTCTG 3'
